# Evaluation of a prototype Orbitrap Astral Zoom mass spectrometer for quantitative proteomics – Beyond identification lists

**DOI:** 10.1101/2025.05.30.657132

**Authors:** Chris Hsu, Nicholas Shulman, Hamish Stewart, Johannes Petzoldt, Anna Pashkova, Deanna L. Plubell, Eduard Denisov, Bernd Hagedorn, Eugen Damoc, Brendan X. MacLean, Philip Remes, Jesse D. Canterbury, Alexander Makarov, Christian Hock, Vlad Zabrouskov, Christine C. Wu, Michael J. MacCoss

## Abstract

Mass spectrometry instrumentation continues to evolve rapidly yet quantifying these advances beyond conventional peptide and protein detections remain challenging. Here, we evaluate a modified Orbitrap Astral Zoom mass spectrometer (MS) prototype and compare its performance to the standard Orbitrap Astral MS. Across a range of acquisition methods and sample inputs, the prototype instrument outperformed the standard Orbitrap Astral MS in precursor and protein identifications, ion beam utilization, and quantitative precision. To enable meaningful cross-platform comparisons, we implemented an ion calibration framework that converts signal intensity from arbitrary units to ion per second. This benchmarking strategy showed that the prototype sampled 30% more ions per peptide than the original Orbitrap Astral MS. This increase in the ion beam utilization resulted in improved sensitivity and quantitative precision. To make these metrics broadly accessible, we added new metrics to the Skyline document grid to report the number of ions measured in a spectrum at the apex of the elution peak or the sum of ions between the peak integration boundaries. Taken together, our results demonstrate the prototype Orbitrap Astral Zoom as a high-performance platform for DIA proteomics and establish a generalizable framework for evaluation of mass spectrometer performance based on the number of ions detected for each analyte. Data are available on Panorama Public and on ProteomeXchange under the identifier PXD064536.

## Introduction

The rapid advancement of mass spectrometry (MS) hardware has exposed a critical need to evaluate improvements in new instruments, such as the Thermo Scientific™ Orbitrap™ Astral™ mass spectrometer (MS)^1–6^. Frequently, new instruments are evaluated based on the number of identified peptides or proteins using established search tools^1–13^. These metrics, while important, remain rooted in qualitative assessments. Additionally, the search protein and peptide search results will be highly dependent on the tool used and will not necessarily capture differences in instrument performance^14^. Quantitative metrics such as transmission efficiency, ion beam utilization, and measurement precision or accuracy are challenging to quantify and compare across instrument platforms. Moreover, new instrument platforms could introduce capabilities that are not immediately compatible with existing data processing workflows, yet they could offer substantial benefits if analytical tools are adapted to fully utilize them.

To enable direct comparisons between different instrument platforms, we would ideally inject a constant amount of sample into two instruments or run them under different settings and then report the number of ions measured by the detector. However, a fundamental challenge lies in the fact that the scale of the reported intensities is often unknown, as they are typically expressed in arbitrary units. While vendors may calibrate detector gain, there remains a need for a universal strategy to convert vendor-specific intensity units into a standardized scale of ions per second. To address this gap, we present a strategy that leverages the relationship between the number of ions measured and the precision of an ion current ratio measurement to derive a correction factor between reported intensity and ions per second^15^. This calibration can be performed using a simple infusion experiment on any instrument platform. We use this calibration factor to illustrate both differences and similarities in the reported signal intensities across various Thermo Scientific analyzers and detection technologies.

To capitalize on this calibration of intensity to ions/second, we added new reporting capabilities to the Skyline^16,17^ document grid to enable extraction of peptide level metrics from liquid chromatography-mass spectrometry data. These metrics include for each peptide 1) the total number of ions in the spectrum at the apex of the chromatographic peak, 2) the number of ions from the extracted precursor > product ion transitions for the target peptide at the chromatographic peak apex, and 3) the number of ions from the target peptide across the chromatographic peak integration boundaries. These metrics can be used to assess both targeted and untargeted measurements.

Using these metrics, we evaluate the effect of modifications to a prototype Orbitrap Astral Zoom mass spectrometer (MS) platform designed to improve instrument efficiency (Figure 1). This modified instrument was improved in three ways. The overhead time from ion pipelining, which contributes to duty cycle loss, was reduced, resulting in an increase in cycle time. This enabled the same ion utilization to be performed in less time. Second, the Orbitrap Astral MS was modified to enable pre-accumulation of ions in the bent square quadrupole region of the ion source prior to the quadrupole mass analyzer as described previously^18^. This pre-accumulation enables an improvement in the effective ion beam utilization by the instrument. Third, this modified instrument made use of improved signal processing that could theoretically deconvolute incompletely resolved fragments in the Astral mass spectra. Using HeLa and extracellular vesicle peptide inputs, we benchmarked both instruments across multiple DIA acquisition settings, assessing acquisition rate, ion counts, precursor and protein detection, and quantitative precision. Our results demonstrate that the Prototype Orbitrap Astral MS consistently achieves faster cycle times, higher ion sampling, and improved detection performance, highlighting its potential for high-throughput and biologically relevant applications.

**Figure 1.**
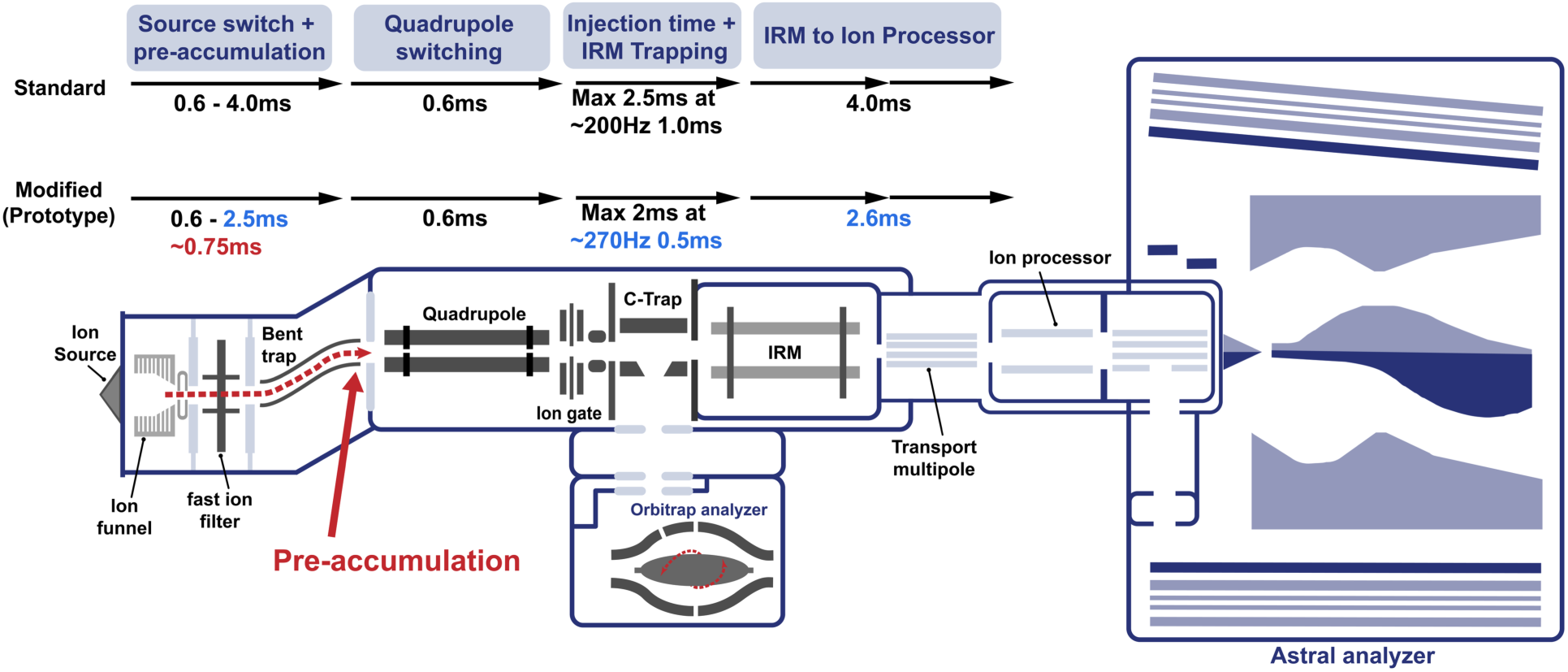
Schematic of the Orbitrap Astral Zoom MS prototype. A newly introduced pre-accumulation step (in red) allows for longer injection times for the Orbitrap Astral Zoom MS prototype. Additional timing optimizations (in blue) show reduced overhead throughout the ion transfer process compared to the standard Orbitrap Astral MS, enabling higher acquisition rates and improved duty cycle efficiency.

## Experimental Methods

### HeLa samples preparation

HeLa cells were grown until 70% confluent and lysed in a buffer containing 2% SDS, 100 mM Tris-HCl (pH 8.5), and Thermo Fisher protease inhibitors, and the lysates were briefly sonicated using a Branson probe sonicator. The total protein concentration was quantified using the bicinchoninic acid (BCA) assay (BCA kit, Thermo Fisher Scientific) using bovine serum albumin standards. The lysate was diluted to 1 µg/µL, reduced with 20 mM dithiothreitol (DTT), and alkylated with 40 mM iodoacetamide (IAA). Protein aggregate capture was used for cleanup protocol^19,20^. Briefly, proteins were bound to ReSyn hydroxyl magnetic beads at 4 µL beads per 25 µg protein by adding acetonitrile to 70% final concentration. The beads were washed with three cycles of 95% ACN and two cycles of 70% ethanol. Residual ethanol was removed by centrifugation. The washed beads were resuspended in 50 mM ammonium bicarbonate containing trypsin (1:20 enzyme-to-protein ratio) and incubated at 47 °C for 3 hours. After digestion, peptides were eluted from the beads and dried using a speedVac vacuum centrifuge. The dried peptides were stored at -80°C until analysis. Prior to LC-MS, the frozen peptides were reconstituted in 0.1% formic acid to a final concentration of 0.2 µg/µL.

### Plasma Preparation and extracellular vesicle (EV) Enrichment

Plasma membrane particle enrichment was performed on a Thermo Scientific KingFisher Flex instrument following the Mag-Net protocol, a magnetic bead approach developed by Wu et al. (2024)^21^. The same protocol was described in Heil et al (2023)^2^. To start, 100 µL of plasma was first mixed with protease and phosphatase inhibitors and then combined with an equal volume of Binding Buffer (BB) containing 100 mM Bis-Tris Propane, pH 6.3, 150 mM NaCl. MagReSyn strong anion exchange beads (ReSyn Biosciences) were pre-equilibrated twice in Equilibration/Wash Buffer (WB) containing 50 mM Bis-Tris Propane, pH 6.5, 150 mM NaCl). These beads were mixed with gentle agitation and then added to the plasma:BB mixture in a 1:4 bead-to-plasma ratio, followed by a 45-minute incubation at room temperature. Post-incubation, the beads were washed three times with WB for five minutes each using gentle agitation. Membrane particles bound to the MagReSyn beads were solubilized by a lysis buffer containing 1% SDS, 50mM Tris at pH 8.5, 10mM TCEP, and 800ng yeast enolase protein. Post reduction, 15mM iodoacetamide was added and incubated for 30 minutes in the dark and quenched with 10mM DTT for 15 minutes. For sample clean-up, protein aggregate capture (PAC) was performed by adding acetonitrile to a final concentration of 70% to precipitate the proteins at room temperature for 10 minutes^19,20^. The beads were washed three times in 95% acetonitrile and two washes in 70% ethanol for 2.5 minutes per wash, using the magnetic field on the KingFisher Flex to sepa rate the beads. After clean-up, the proteins were digested at 47°C for 1 hour in a 20:1 ratio of trypsin-to-protein containing 100 mM ammonium bicarbonate. Digestion was halted by adding 0.5% formic acid, and an internal control, Pierce Retention Time Calibrant peptide cocktail (Thermo Fisher Scientific), was included at a final concentration of 50 fmol/uL. The peptides were dried using a speedVac vacuum centrifuge and frozen until analysis.

### Evaluation of acquisition rate and cycle time

Data-independent acquisition (DIA) was used to assess the theoretical limit of acquisition rate and cycle time for the Orbitrap Astral Zoom MS prototype and Orbitrap Astral MS. The DIA method consisted of a single MS1 scan using the Orbitrap detector at 240,000 resolving power, precursor range of 375-985 *m/z*, 50ms maximum injection time, “standard” automatic gain control (AGC), and loop time set to 1000 seconds. After the only MS1 spectrum was collected, MS2 spectra were acquired continuously for 1 minute using the Astral analyzer on the Orbitrap Astral Zoom MS prototype. This prototype instrument could also operate as a standard Orbitrap Astral MS with the overhead timing and standard Astral spectrum processing. The MS2 spectra included the following settings: 2 Th or 4 Th isolation windows, 30% HCD collision energy, 100% AGC target, and a precursor mass range of 400–900 m/z using the optimized window placement^22^. The maximum injection time varied from 0.5 - 10 milliseconds (ms). The Thermo Fisher H-ESI source was used with the spray voltage set to 0 V to force the instrument to always reach the maximum injection time.

Post-acquisition, the raw files were converted to mzML using MSConvert (version 3.0.25073) in ProteoWizard and processed in R using the mzR package^29^. The acquisition rate was calculated as the total number of spectra divided by total acquisition time (or total retention time). The overhead was calculated as the difference between the average time per MS2 scan subtracted by the set injection time in the method. And the cycle time was calculated as the retention time difference between one full cycle of precursor scan of 400-900 *m/z*.

### Liquid-chromatography and mass spectrometry

HeLa and extracellular vesicle (EV) samples were analyzed using a Thermo Scientific Vanquish Neo UHPLC system coupled to an Orbitrap Astral Zoom MS prototype. This prototype instrument could operate as a commercially available Orbitrap Astral MS with the overhead timing and standard Astral spectrum processing (referred to as Orbitrap Astral MS). This instrument could also be operated with the reduced ion pipelining overhead, pre-accumulation, and Astral spectral deconvolution (referred to as Prototype). Peptides were separated via a 24-minute gradient at 1.3 µL/min, with the following stepwise elution profile: 4–6% Buffer B over 0.7 minutes, 6–6.5% Buffer B over 0.3 minutes, 6.5–40% Buffer B over 20 minutes, 40–55% Buffer B over 0.5 minutes, and 55–99% Buffer B over 3.5 minutes for column washing. Following separation, data-independent acquisition (DIA) was performed using a cycle comprising a full MS1 acquisition on the Orbitrap detector at 240,000 resolving power, precursor range of 375–985 *m/z*, 50 ms injection time, and standard automatic gain control (AGC). Following each MS1, MS2 spectra was collected using the Astral detector with the following parameters: 3 Th non-staggered isolation windows, 6 ms injection time, precursor mass range of 400–900 *m/z*, 27% higher-energy collisional dissociation (HCD) collision energy, and 200% AGC target. For isolation window optimization, three configurations were tested while maintaining consistent cycle times: 2 Th isolation windows with 4 ms injection time, 3 Th windows with 6 ms injection time, and 4 Th windows with 8 ms injection time. For the Orbitrap Astral MS, the same configurations were used with the same LC and column setup. The expected chromatographic peak width was set to 6 seconds for all sample runs.

### DIA data processing

An *in-silico* spectral library was generated using Carafe^30^ by processing a representative HeLa sample run. Following library generation, all data files were searched with DIA-NN (v2.1.0). For peptide identification, the precursor ions were modeled with the following parameters: trypsin protease specificity with 1 missed cleavage allowed, fixed carbamidomethylation of cysteines, and precursor charge range of 2+ to 3+. The resulting spectral library was exported in a format compatible with Skyline for downstream analysis. The settings of the DIA-NN algorithm included match-between-runs (MBR), “Protein inference”, and “Unrelated Runs”. Machine learning-based scoring was enabled using “NNs (cross-validated)”, Cross-run normalization was set to “Global”, and library generation utilized “IDs profiling”.

### Instrument calibration to ions/sec

#### Theory and derivation

To enable accurate and meaningful comparisons between different mass spectrometers, it is necessary to calibrate the instruments such that signal intensity measurements are expressed on a common scale—typically in ions per second (ions/sec). This calibration is crucial for benchmarking instrument sensitivity and noise characteristics, especially when comparing systems with different hardware configurations (e.g., analyzers, ion optics) or acquisition strategies.

To perform this calibration, we carry out repeated measurements on a consistent peptide standard or analyte compound to assess signal precision, where noise is defined as the variability observed when measuring a stable signal. The variability associated with these repeated measurements is the total variance (σ^2^**T**). The total variance originates from two primary sources: Poisson (shot) noise (σ^2^**P**) and additional sources of variation (σ^2^**O**), such as detector electronics, signal processing, and digitization artifacts. This is adapted directed from Canterbury et al.^15^ as Equation (1):

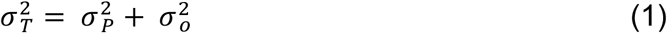

To isolate and quantify these noise components, we acquire many nominally identical tandem mass spectra (typically 5000) from a stable analyte or peptide standard. From each spectrum, we extract the intensities of the same set of prominent fragment ions. Rather than analyzing absolute intensities, we compute intensity ratios between a set reference fragment ion and each of the other selected prominent fragment ions across spectra. This ratio metric approach helps to cancel out common-mode noise sources, such as fluctuations in electrospray stability or overall ionization efficiency, which would otherwise affect all peaks proportionally.

To relate measured signal intensity to the actual number of ions detected, we consider the basic relationship between these quantities. Equation (2) models the relationship between signal intensity and number of ions. The measured intensity *I* of any given signal in a spectrum is proportional to the number of ions *N* divided by the time it takes to accumulate ions in the trap or injection time *t*. The α represents a variable that accounts for instrument-specific scaling (e.g., differences in detector gain, analyzer).

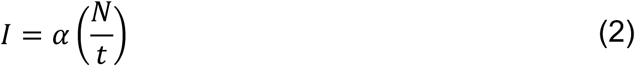

Because both α and *t* are constants, the ratio (R) between two given intensities can be defined as the ratio between two reported *N* values in Equation (3). The assumption is that the two intensities, *I*ₐ and *I*b, are measured under the same α and *t* conditions. Therefore, these constants cancel out for the same measurements.

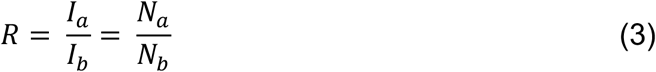

Propagation of errors can be used to derive an equation for how the variance in the ratio (σ^2^ᵣ) is affected by the variance of individual measurements, R itself, and the number of ions that contribute R (Nₐ and Nb), as shown in Equation (4):

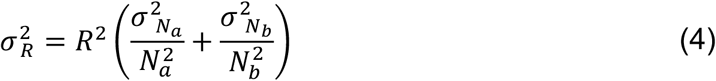

Ion counting follows Poisson statistics, where the variance is proportional to the number of ions. The precision in the ratio from Equation (4) can be further simplified by inserting the variance from Equation (5). The ion count has a direct relationship with the precision: as Nₐ and Nb increase, the variance increases. However, the signal increases more than the noise.

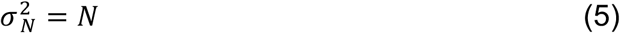

Using the relationship between variance and the number of ions from Poisson counting statistics^25^, Equation (4) is re-written as:

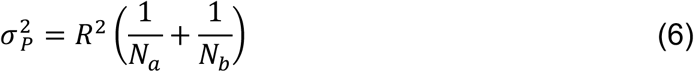

Using Equation (2), we can rewrite Equation (5) in terms of the measured intensity *I* and the injection or dwell time *t*. The product of *I and t* is an uncalibrated estimate of the number of ions and α is a correction factor to convert between uncalibrated ions and calibrated ions.

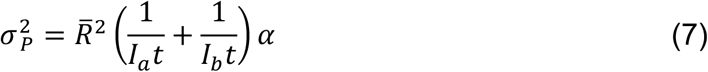

We are using the average of the R and the uncalibrated ions from many measurements as an estimate of the actual ratio and ions. The total variance σ^2^T is the experimentally measured variance from the measurement of the ion current ratio and should include the variance from Poisson noise σ^2^P and noise sources independent of the signal intensity, σ^2^O, as shown in Equation (1). Substituting the variance from Poisson noise from Equation (7) gives us Equation (8).

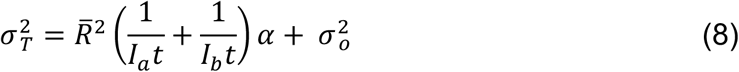

Equation (8) closely resembles the form of a linear equation, *y = mx + b*, where the total variance σ^2^T corresponds to the dependent variable *y*, the slope is defined by α, the independent variable *x* is a function of the measured signal intensity *I* and injection time *t*, and the intercept σ^2^ₒ represents other non-Poisson sources of variation. This final equation allows for an empirical estimation of both the instrument-specific α scaling factor and noise contributions^15,23–25^.

#### Calibrating Instrument Response

Prior to calibrating the instrument response factor, instruments were calibrated using the standard auto-calibration routines using Pierce™ FlexMix™ (Thermo Fisher A39239) as recommended by the manufacturer. The Glu[1]-Fibrinopeptide B peptide (Millipore sigma F3261) was resuspended in a mixture of methanol, water, and formic acid at a ratio of 50:49.9:0.1 and diluted to a final concentration of 1 µM as previously described by Canterbury, et. al^16^. The peptide was infused using a syringe pump with a flow rate of 4 µL/min, and the spray stability was monitored for two minutes prior to data collection. Data were subsequently acquired for instruments using the Astral, the ion trap, and Orbitrap analyzers with the following instruments: Orbitrap Astral Zoom MS prototype, standard Orbitrap Astral MS, Orbitrap Ascend MS, Exploris 480 MS, Stellar MS, and Orbitrap Fusion

Lumos Tribrid MS. For all instruments and analyzers, the precursor isolation was set to m/z ∼786 *m/z* at the 2+ charge state, and 5000 MS2 spectra were collected with 2 *m/z* isolation windows, 100% AGC target, and “auto” injection time. For Orbitrap analyzers, the resolution was set to 30,000 resolving power, and ion traps were set to “Rapid”. The raw MS2 data were converted to mzML files by MSConvert (version 3.0.22175) for downstream analysis. Fragment ions used for Glu[1]-Fibrinopeptide B peptide instrument calibration included *m/z* 480.23, 627.32, 684.35, 813.39, 942.43, 1056.47, 1171.50, and 1285.54 with the *m/z* 942.4291 fragment selected as the reference ion for the linear regression.

For Ultramark 1122, MS2 acquisition was performed using similar instrument settings as described above, with a normalized collision energy of 48% and a range of 200–1200 m/z. Fragment ions used for calibration included m/z 553.94, 665.96, 677.98, 765.95, 777.97, 789.99, 877.96, 889.98, and 989.97, with m/z 989.9734 selected as the reference ion for calibration. For Ultramark 1822, the normalized collision energy was set to 47%, and the mass-to-charge range was extended to 200– 2000 m/z to capture higher-mass fragments. Calibration was performed using fragment ions at m/z 753.93, 853.92, 945.93, 965.95, 1065.93, 1177.94, 1277.94, 1377.93, 1489.94, 1589.94, and 1689.94, with m/z 1377.93 designated as the reference ion.

## Results

### Modifications to the Orbitrap Astral MS improve the MS/MS acquisition rate

To assess performance differences between the Orbitrap Astral MS and the Orbitrap Astral Zoom MS prototype, we compared acquisition rate, acquisition overhead, and cycle time across varying injection times and DIA isolation window configurations. To ensure that the MS2 acquisitions reached their maximum allowable fill times, we set the H-ESI source to 0 V, disabling electrospray ionization and forcing the system to accumulate ions until the maximum injection time was reached. DIA data were acquired across a range of injection times using three isolation widths: 2 Th, 3 Th, and 4 Th.

Acquisition rate, calculated as the total number of spectra divided by total acquisition time, was consistently higher on the Orbitrap Astral Zoom MS prototype across all conditions (Figure 2A). The largest differences were observed at shorter injection times (1–3 ms). Under the fastest conditions (2 Th with 4 ms injection time), the Orbitrap Astral Zoom MS prototype approached the theoretical acquisition ceiling of 270 Hz, while the Orbitrap Astral MS rate peaked around ∼217 Hz. The Orbitrap Astral MS also showed a larger decline in acquisition rate when using wider isolation windows, likely due to increased acquisition transition costs. To quantify the source of these differences, we calculated acquisition overhead as the difference between MS2 acquisition duration and the set injection time (Figure 2B). The Orbitrap Astral Zoom MS prototype consistently exhibited ∼1 ms lower overhead compared to the Orbitrap Astral MS across all injection times, and although overhead increased slightly with wider isolation windows, the instrument remained more efficient under all conditions, with minimal performance penalty. These improvements directly translated into shorter duty cycles (Figure 2C), with the Orbitrap Astral Zoom MS prototype consistently achieving faster cycle times across the full 400–900 m/z precursor range.

**Figure 2.**
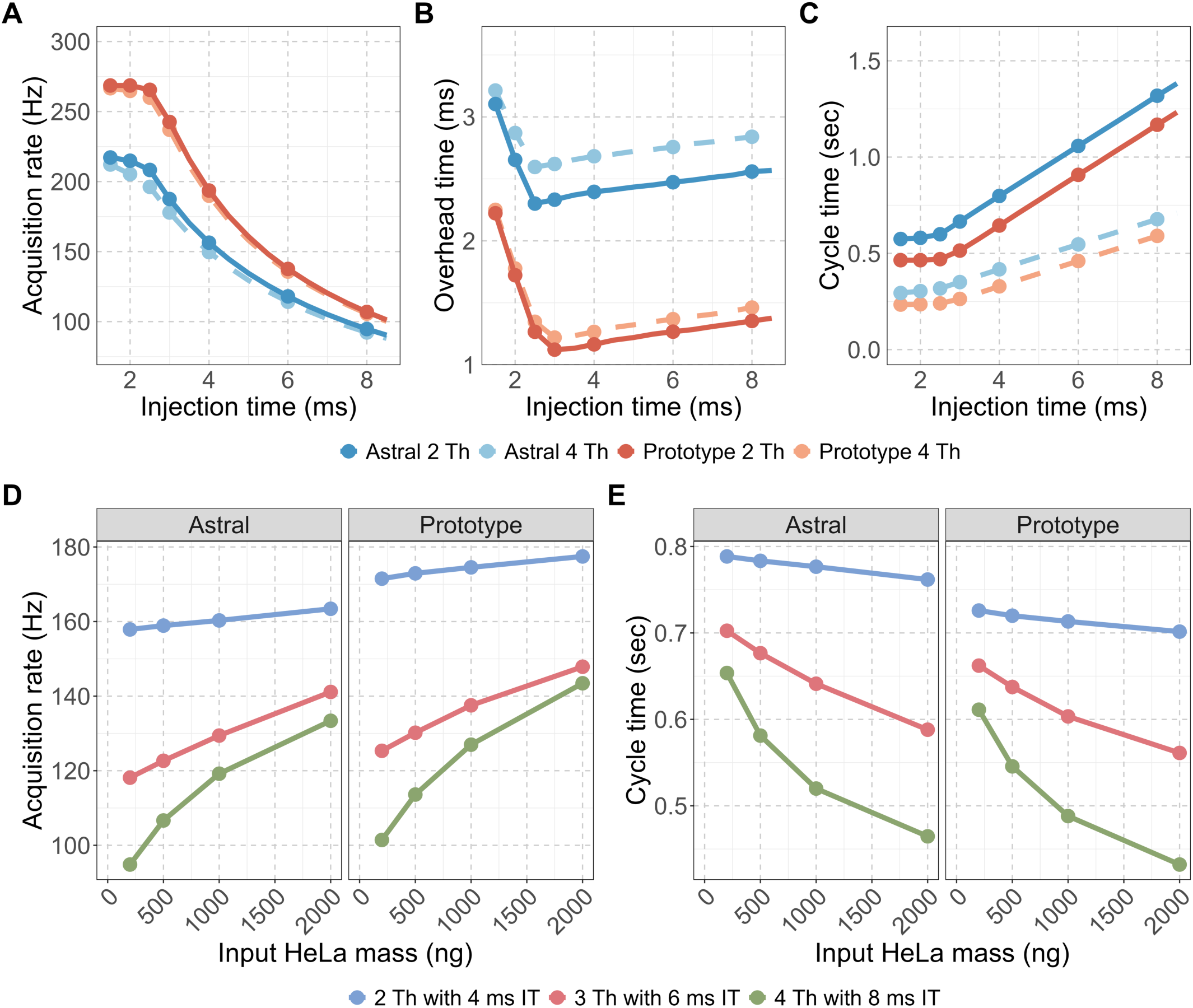
The Orbitrap Astral Zoom MS prototype reduces acquisition overhead and improves MS2 acquisition rate and cycle time. Evaluation of two DIA configurations: narrow (2 Th) and wider (4 Th) isolation windows for (A) MS2 acquisition rate (Hz), (B) Overhead time, calculated as the difference between the mean MS2 acquisition time and injection time, and (C) Cycle time for a 400-900 m/z precursor range. Panels (D-E) show the impact of instrument and DIA method on performance across increasing HeLa peptide input for (D) Acquisition rate and (E) Cycle time are shown for three DIA configurations with set injection times (IT): 2 Th with 4 ms IT (blue), 3 Th with 6 ms IT (red), and 4 Th with 8 ms IT (green).

We next evaluated how these improvements translated into practical performance using biological samples. The acquisition rate and cycle time were measured across a range of HeLa peptide inputs (200–2000 ng) using the three different DIA methods: 2 Th with 4 ms injection time (IT), 3 Th with 6 ms IT, and 4 Th with 8 ms IT. At all input levels and DIA methods, the Orbitrap Astral Zoom MS prototype demonstrated improved acquisition rate and decreased cycle time over the Orbitrap Astral MS. These improvements were most pronounced at higher sample loads, where ion availability enabled automatic gain control (AGC) targets to be met faster, allowing the full benefit of the Prototype’s faster acquisition transitions and reduced overhead to be utilized. Impressively, the Orbitrap Astral Zoom MS prototype achieved these performance benchmarks with a slightly longer injection time of ∼0.75 ms over the standard Orbitrap Astral MS (Figure 1, Figure S1) with the addition of the pre-accumulation step. These further benchmark the faster performance and efficiency of this new instrument.

### The Orbitrap Astral Zoom MS prototype improves peptide and protein detections and quantitative precision for DIA

Across all conditions and different HeLa loading amounts, the Orbitrap Astral Zoom MS prototype detected more precursors and proteins than the standard Orbitrap Astral MS (Figure 3A–B). These differences were particularly pronounced at higher peptide loads (≥500 ng), where statistical testing showed p-values < 0.05 in both precursor and protein identifications (Tables S1–S2). For example, at 2000 ng and 3 Th with 6 ms maximum injection time (maxIT), the Prototype yielded on average 231,842 ± 365 detected precursors and 9784 ± 22 proteins, compared to 216,226 ± 207 precursors and 9677 ± 15 proteins on the Orbitrap Astral MS (p = 0.001 and 0.024, respectively). The most pronounced absolute gains in precursor identifications were observed with the widest isolation window (4 Th with 8 ms maxIT), where the Prototype instrument was able to detect 224,061 ± 173 precursors at 2000 ng input (vs. 204,826 ± 199 on Orbitrap Astral MS, p < 0.001) at a false discovery rate of less than 0.01 using DIA-NN v2.1.0.

**Figure 3.**
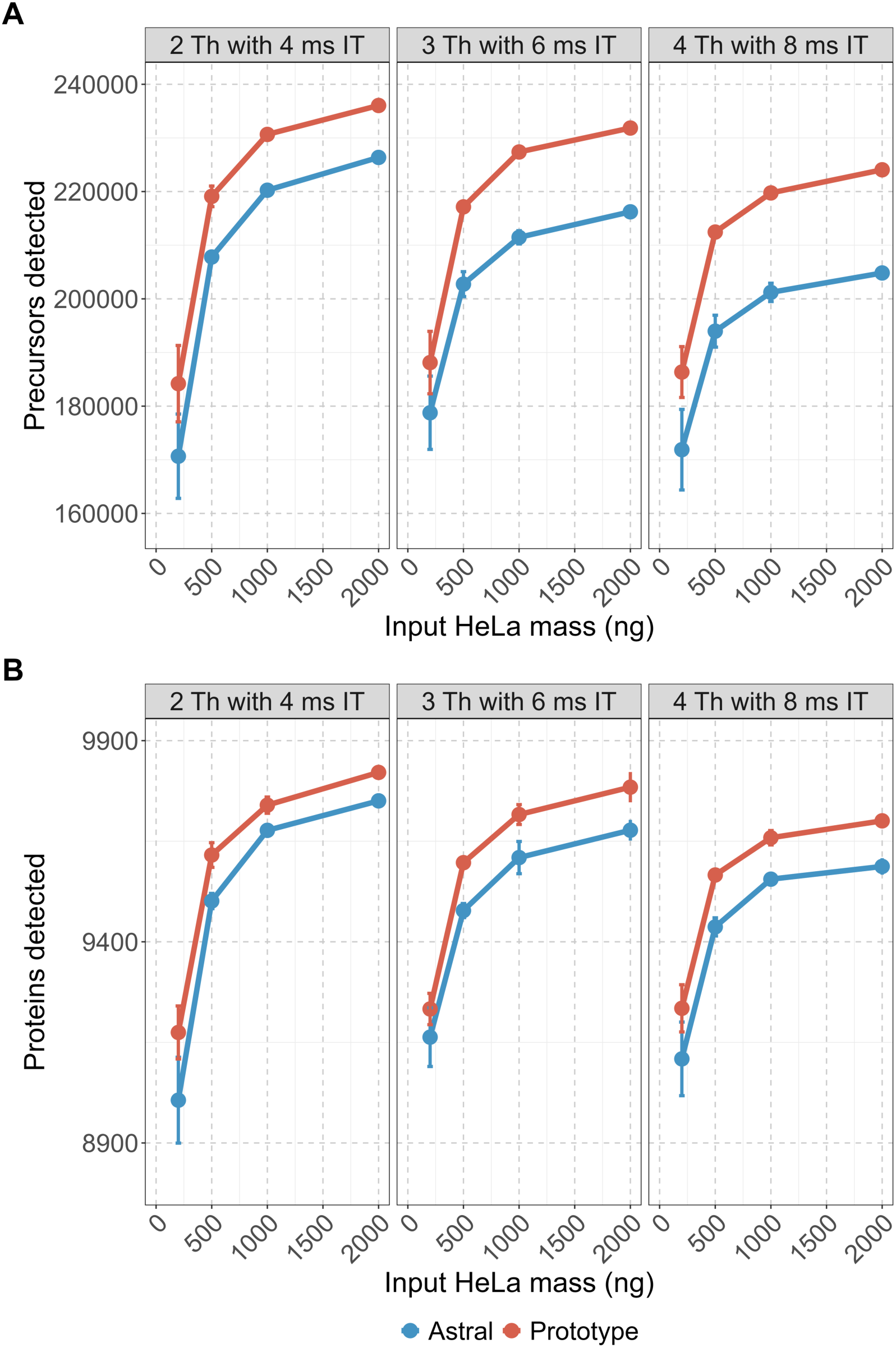
Orbitrap Astral Zoom MS prototype improves precursor and protein detection across different DIA isolation windows for higher input peptide loads. HeLa peptide digests ranging from 200, 500, 1000, and 2000 ng were analyzed in triplicate using three DIA acquisition schemes: 2 Th with 4 ms, 3 Th with 6 ms, and 4 Th with 8 ms MS2 injection times (IT). All experiments were performed under matched LC conditions on both instrument platforms. (A) Average precursor and (B) protein detections reported by DIA-NN are shown across various HeLa peptide load amounts for the Orbitrap Astral Zoom MS prototype (red) and Orbitrap Astral MS (blue) with included error bars.

The Orbitrap Astral Zoom MS prototype also retained substantial sensitivity at lower HeLa peptide inputs. Although this analysis does not reflect a direct pairwise comparison of identified precursors or proteins across input amounts, the overall coverage at 200 ng remained high relative to 2000 ng input level. For instance, under the 2 Th with 4 ms maxIT condition, the Orbitrap Astral Zoom MS prototype detected 184,193 ± 4109 precursors and 9175 ± 38 proteins at 200 ng. This represents ∼78% and ∼94%, respectively, of the total number of precursors and proteins identified at 2000 ng. The difference reflects a relatively small drop in proteome coverage despite a 10-fold decrease in input mass. The narrowest isolation window (2 Th) yielded the highest number of detections overall, consistent with expectations based on improved specificity. Together, these results underscore the importance of high acquisition speed and optimized ion transmission in maintaining deep proteome coverage across a broad dynamic range of sample inputs.

To further evaluate the quantitative reproducibility and precision of each instrument, we examined the coefficient of variation (CV) for peptide- and protein-level abundance (Figure 4A–B). Both instruments demonstrated strong quantitative reproducibility and precision. The peptide-level median CVs were consistently below 20%, and protein-level median CVs were below 10%. At nearly all inputs (≥500 ng), the Orbitrap Astral Zoom MS prototype exhibited narrower CV distributions and lower median CVs than the Orbitrap Astral MS. These results highlight that both platforms are well-suited for precise DIA-based quantification, but overall, the Orbitrap Astral Zoom MS prototype has better precision and reproducibility in measurements.

**Figure 4.**
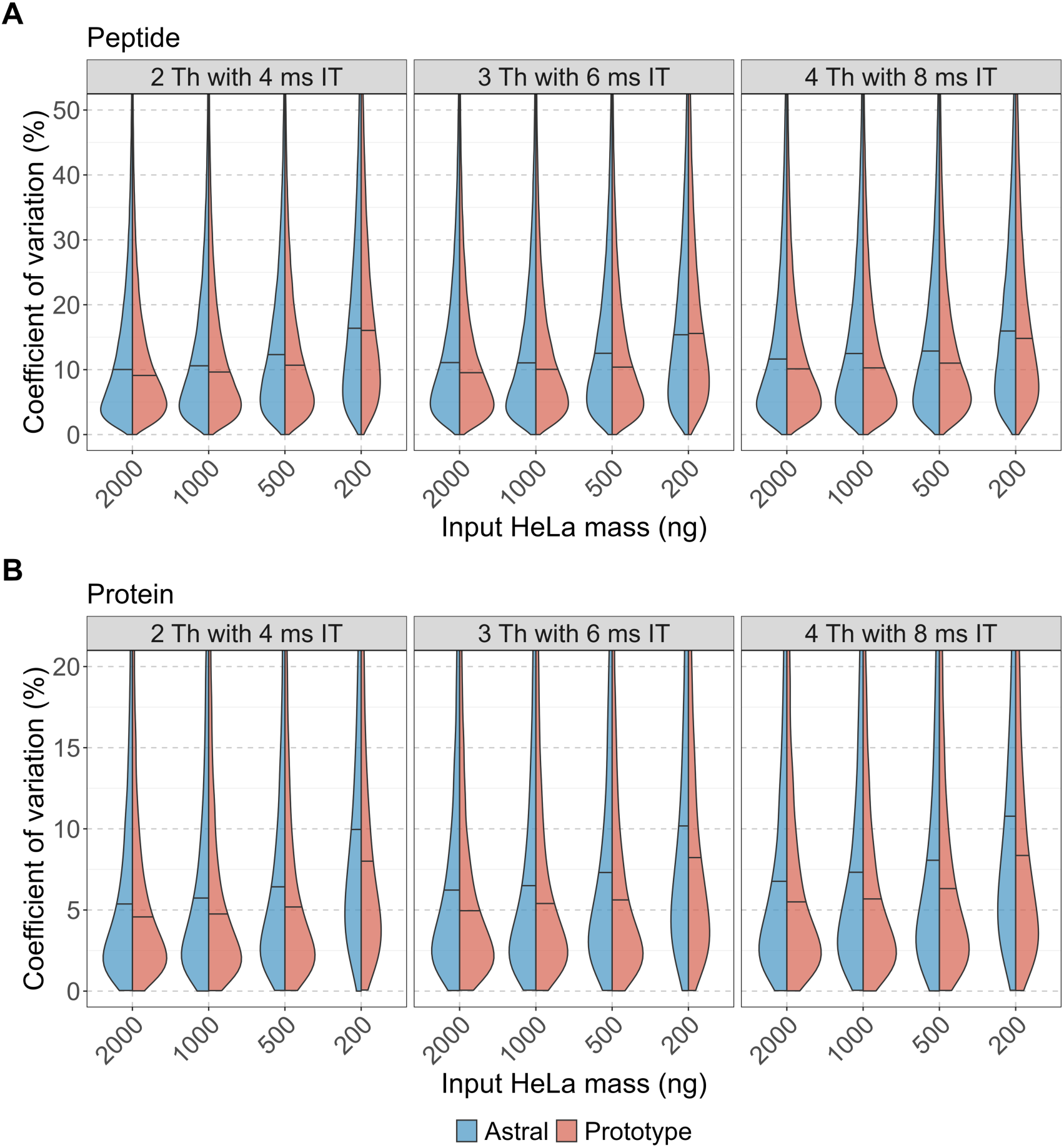
Quantitative precision of peptide and protein measurements across input amounts and DIA acquisition schemes. Coefficient of variation (CV) distributions are shown for **(A)** peptides and **(B)** proteins identified across triplicate DIA runs of a HeLa digest on the Orbitrap Astral MS (blue) and Orbitrap Astral Zoom MS prototype (red). CVs were calculated from median-normalized intensities for each analyte across five input amounts (200–2000 ng) and three DIA acquisition settings (2 Th with 4 ms injection time, 3 Th with 6 ms, and 4 Th with 8 ms). Horizontal lines within each split violin represent the median CV for that condition.

**Figure 5.**
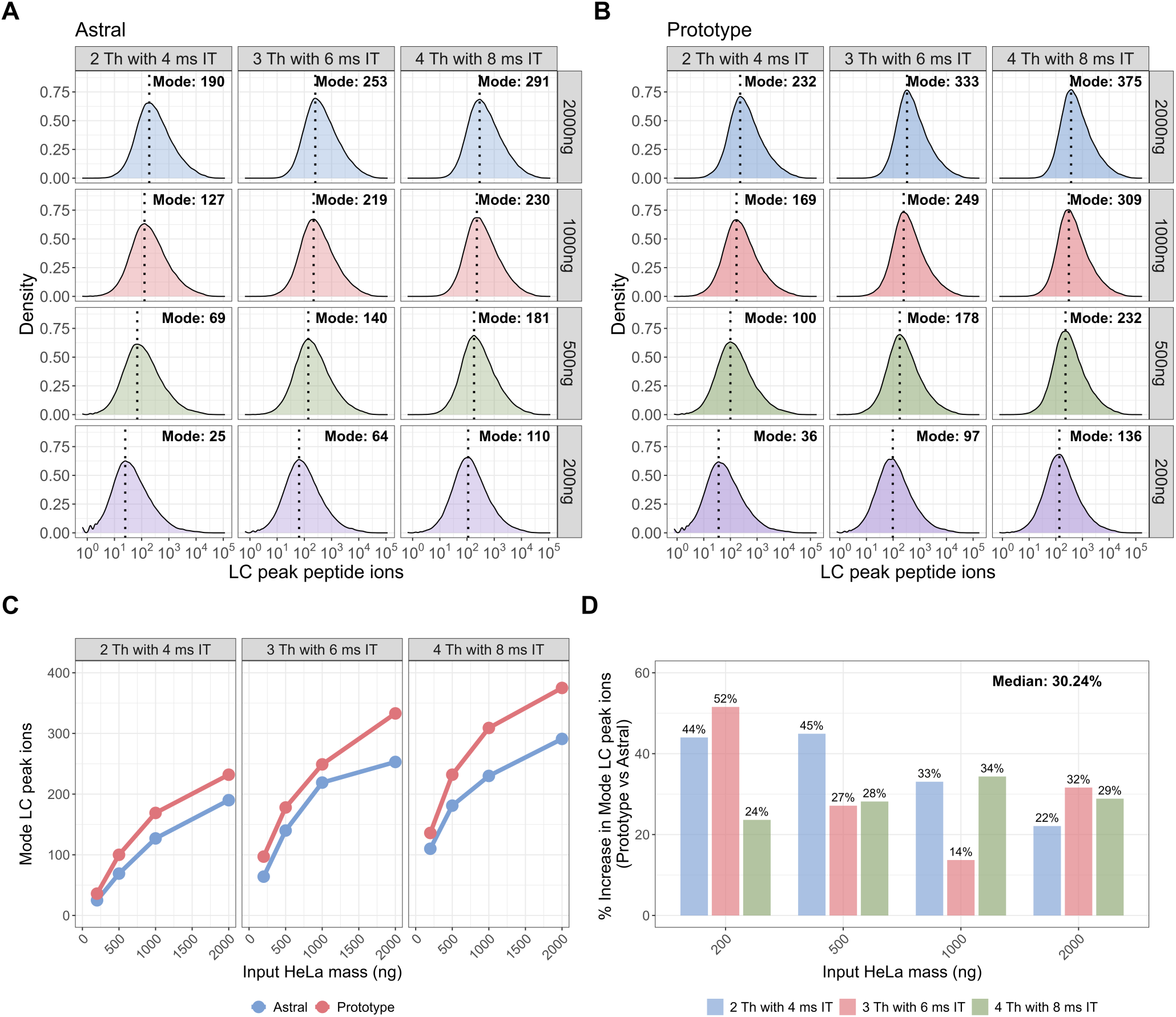
The Orbitrap Astral Zoom MS prototype improves LC peak peptide ion counts across sample inputs and DIA methods. Density plots of LC peak peptide ion counts across various HeLa inputs (200-2000ng) for each DIA method (2 Th, 3 Th, 4 Th) are shown separately for the Orbitrap Astral MS **(A)** and Orbitrap Astral Zoom MS prototype **(B)**. Each panel displays the distribution of calibrated peptide ion counts across all identified peptides. Vertical dotted lines indicate the mode of each distribution. **(C)** Line plot showing the mode of LC peak peptide ion for the same HeLa peptide inputs and DIA isolation window settings comparing the Orbitrap Astral MS (red) and the Orbitrap Astral Zoom MS prototype (blue) instrument with the **(D)** percent increase in mode LC peak peptide ion counts with the reported median percent increases across all conditions.

### Instrument calibration to ions per second

The Orbitrap Astral Zoom MS prototype demonstrated lower CVs in peptide and protein quantification compared to the standard Orbitrap Astral MS. Because CVs will improve with an increase in the number of ions measured^25^, this observation raised the question of whether the improved precision could be attributed to a greater number of ions measured per peptide. To enable meaningful comparisons between different mass spectrometers, it was necessary to calibrate the instruments to the same signal intensity measurements on the same scale – typically reported in ions per second (ions/sec).

To standardize the signal intensity across instrument platforms to ions/sec, we applied an ion calibration method based on Poisson-based ion statistics (see Equation 8 and Methods). This calibration framework derives a correction factor (α) by regressing the total variance from fragment intensity ratios against the inverse of the estimated ion counts across repeated measurements. To visualize the steps in this process, we analyzed the MS/MS spectra of Glu[1] Fibrinopeptide B, a standard peptide for instrument tuning and calibration (Figure S2A). First, the raw signal intensities of several prominent fragment ions were plotted and then ion counts were computed with the injection time (Figure S3A-B). The intensity ratios of each fragment ion to a reference fragment ion, 942.4291 *m/z*, were computed (Figure S3C). Linear regression plots were constructed by showing the intensity variance against the number of ions and the intensity ratios of two fragments. From these plots, we extracted linear equation containing the α (slope), σ₀^2^ (y-intercept), and R^2^ (goodness-of-fit) values. Table 1 summarizes calibration results for Glu[1]-Fibrinopeptide B across five Thermo Scientific mass spectrometers spanning three analyzer types: Astral, Orbitrap, and ion trap. An α value of 1.0 indicates that signal intensities are already accurately scaled to ions/sec. Alpha values greater than 1.0 suggest an overestimation of ion counts (i.e., reported intensity is higher than true ion current), while values less than 1.0 suggest underestimation.

**Table 1.**
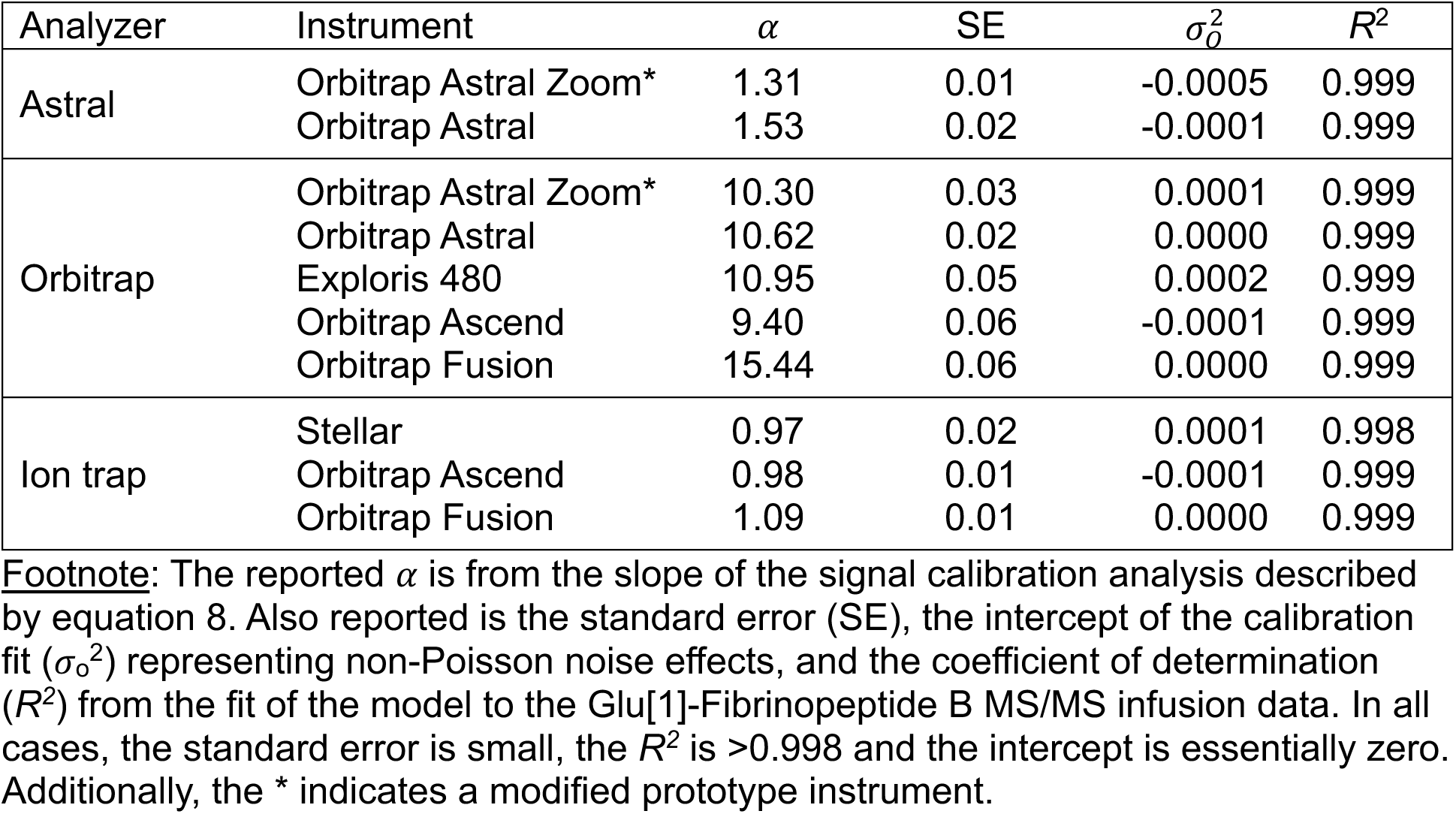
Instrument calibration results for Glu[1]-Fibrinopeptide B.

| Analyzer | Instrument | $\alpha$ | SE | $\sigma_o^2$ | $R^2$ |
| --- | --- | --- | --- | --- | --- |
| Astral | Orbitrap Astral Zoom* | 1.31 | 0.01 | -0.0005 | 0.999 |
|  | Orbitrap Astral | 1.53 | 0.02 | -0.0001 | 0.999 |
| Orbitrap | Orbitrap Astral Zoom* | 10.30 | 0.03 | 0.0001 | 0.999 |
|  | Orbitrap Astral | 10.62 | 0.02 | 0.0000 | 0.999 |
|  | Exploris 480 | 10.95 | 0.05 | 0.0002 | 0.999 |
|  | Orbitrap Ascend | 9.40 | 0.06 | -0.0001 | 0.999 |
|  | Orbitrap Fusion | 15.44 | 0.06 | 0.0000 | 0.999 |
| Ion trap | Stellar | 0.97 | 0.02 | 0.0001 | 0.998 |
|  | Orbitrap Ascend | 0.98 | 0.01 | -0.0001 | 0.999 |
|  | Orbitrap Fusion | 1.09 | 0.01 | 0.0000 | 0.999 |

**Footnote:** The reported $\alpha$ is from the slope of the signal calibration analysis described by equation 8. Also reported is the standard error (SE), the intercept of the calibration fit ( $\sigma_o^2$ ) representing non-Poisson noise effects, and the coefficient of determination ( $R^2$ ) from the fit of the model to the Glu[1]-Fibrinopeptide B MS/MS infusion data. In all cases, the standard error is small, the $R^2$ is $>0.998$ and the intercept is essentially zero. Additionally, the \* indicates a modified prototype instrument.

The instruments associated with Astral and ion trap analyzers exhibited alpha factors very close to 1, which indicates that the instruments are reporting intensities that are well calibrated to ions/sec. The Astral analyzer’s detector is normally internally calibrated to the areas of defocused single ion peaks, measured at heightened detector gain, and signal intensity reported in ions per second. As a comparative test, an Orbitrap Astral MS instrument was set to a similar defocused condition with heightened detector gain, and the number of single ion peaks detected per scan matched within 10% the number of ions predicted by the calibration^27^. The intensity measured by mass spectrometers with Orbitrap mass analyzers (e.g., Orbitrap Fusion MS, Orbitrap Ascend MS, Exploris 480 MS) need to be divided by 9.4 - 15.44 to obtain the calibrated ions/sec. Interestingly, the Orbitrap Exploris 480 MS, Orbitrap Astral MS, and Orbitrap Ascend MS share the same internal Orbitrap analyzer, but only the Orbitrap Ascend MS results differed, which is most likely due to differences in software. The Orbitrap Fusion Lumos Tribrid MS, being the oldest model, gave the highest calibration factor of 15.44.

To evaluate whether calibration is compound-specific, we extended the analysis to two additional standards: Ultramark 1122 and Ultramark 1822 from Pierce FlexMix calibration solution. These analytes are regularly used for diagnostics and mass calibration across different instrument platforms, which makes it readily available for others to adopt this approach without requiring additional compound purchases or custom preparations. An example MS/MS spectrum of three analyte compounds with key fragments from y- and b-ions for Glu[1]-Fibrinopeptide B (Figure S2A) and major peaks from Ultramark 1122 and Ultramark 1822 (Figure S2B-C). Linear regression results for both Orbitrap Astral MS and Orbitrap Astral Zoom MS prototype instruments using Ultramark compounds demonstrated consistent fits (R^2^ > 0.99) and similar α values across compounds (Figure S4). While α varied slightly by analyte, these differences could be attributed to different *m/z* of the fragments, different range of ratios measured, the range of variance from the measurements.

### Assessing ion counts between Orbitrap Astral Zoom MS prototype and Orbitrap Astral MS

To quantify the number of ions across MS2 spectra, we implemented a new set of custom metrics in the Skyline document grid that enable ion count calculations for any analyte measured with LC-MS data. As summarized in Table S3, these metrics include LC Peak Total Ion Current Area Fragment, Apex Total Ion Count Fragment, LC Peak Total Ion Count Fragment, Apex Analyte Ion Count Fragment, and LC Peak Analyte Ion Count Fragment (also referred to as LC Peak Peptide Ion Count).

For the purposes of this study, we focused on two key metrics: Apex Total Ion Count Fragment and LC Peak Analyte Ion Count Fragment. The apex value is calculated as the product of the total ion current (TIC) and the injection time for the MS2 spectrum containing the highest transition intensity. Notably, this value directly mirrors the ion count reported in the raw file at that retention time. The LC peak peptide Ion Count is computed as the sum of the products of transition intensities and injection times across all MS2 spectra within the peak integration boundaries. This metric approximates the total number of ions contributing to the quantification of a given peptide. Taken together, these metrics establish a quantitative framework for comparing ion counts across instruments, acquisition settings, and peptide intensities. When combined with the appropriate instrument-specific α (alpha) correction factor, they allow for calibrated cross-platform comparisons in units of ions per second.

The Orbitrap Astral MS uses automatic gain control (AGC) to regulate the number of ions used in each spectrum. The injection time is adjusted to maintain a target number of ions in each spectrum. To evaluate whether the data collected accurately reflects the AGC target, we examined the apex total ion count per peptide from different amounts of HeLa peptide digests and the three DIA method settings described previously (Figure S5). Both the Orbitrap Astral MS and the Orbitrap Astral Zoom MS prototype were configured with an AGC target of 20,000 ions, and ion counts were calibrated to ions/sec using instrument-specific α factors (1.53 for Orbitrap Astral MS and 1.31 for Orbitrap Astral Zoom MS prototype). The median ion counts from both instruments reach the AGC target at 500 ng input, particularly for the 3 Th with 6 ms maxIT and 4 Th with 8 ms maxIT settings, suggesting efficient and accurate ion regulation at this input mass. At lower inputs (200 ng), underfilling becomes evident—especially for the narrowest 2 Th window with 4 ms maxIT—where median ion counts fall below the AGC target as the 4 ms maxIT is frequently insufficient to reach 20,000 ions. While the 2 Th and 4 ms maxIT methods might obtain the largest number of protein and peptide detections, it underfills the AGC target most often except at ≥1 µg loading.

To assess differences in peptide-level ion count, we compared LC peak peptide ion counts between the Orbitrap Astral MS and Orbitrap Astral Zoom MS prototype across the same dataset. Density plots of the LC peak ion distributions revealed consistent rightward shifts for the Orbitrap Astral Zoom MS prototype across all conditions (Figure 4A–B), indicating higher peptide-level ion counts. We extracted the mode of each distribution to quantify the most common ion count under each input mass and DIA setting. Across all input levels and DIA configurations, the Orbitrap Astral Zoom MS prototype yielded higher mode LC peak ion counts than the Orbitrap Astral MS (Figure 4C–D). The largest gains were observed at 200 ng, where the Orbitrap Astral Zoom MS prototype delivered up to 52% more ions under the 3 Th with 6 ms maxIT setting (Figure 4E). Even at higher inputs (≥1000 ng), the Prototype maintained a 22–33% increase in the mode number of ions per peptide. To further evaluate performance at the individual peptide level, we performed pairwise comparisons of shared peptides, computing the log₂ fold-change in LC peak ion counts between instruments. These comparisons showed a consistent positive shift favoring the Prototype across all conditions (Figure S6). The greatest increase in log₂ ion ratios were observed in the low-to-intermediate ion count range (approximately 10s to 100s of ions), where the Prototype likely improves sampling of peptides near the detection limit. In contrast, peptides with high ion abundance (e.g., >1000 ions) already produce strong signals on both instruments, leaving less room for improvement.

#### Assessing quantitative performance with human EVs using matrix-matched calibration curve

To benchmark performance differences on a distinct sample type, we evaluated the Orbitrap Astral MS and Orbitrap Astral Zoom MS prototype using a matrix-matched calibration curve^27^ consisting of human extracellular vesicle (EV) peptides diluted into chicken plasma^28^. The EV data was generated using the 3 Th with 6 ms maxIT DIA method on both instruments. We selected this method because it strikes a balance between the 2 Th and 4 Th methods in terms of detections and ion counts. Although the 3 Th method resulted in slightly fewer precursors and protein detections than the 2 Th method, it compensated by enabling more time for the AGC to reach its target, resulting in higher ion counts per peptide (Figure S5). Data was generated for 14 concentrations ranging from 0.1%, 100%, including a 0% Human EV as a control.

Using the 100% human EV condition from this series, we compared peptide and protein identifications, ion sampling, and quantification precision. Consistent with trends observed in earlier datasets (Figure 3–4), the Orbitrap Astral Zoom MS prototype identified more EV precursors and proteins than the Orbitrap Astral MS, while also presenting lower median coefficients of variation (Figure S7A–D). Additionally, protein rank abundance analysis based on the summed LC peak peptide ion counts for the peptides of the same protein revealed that the Orbitrap Astral Zoom MS prototype delivered higher ion counts across a broader dynamic range, particularly for EV-specific marker proteins (Figure S7E).

To assess the quantitative accuracy of the Orbitrap Astral MS and Orbitrap Astral Zoom MS prototype across a dynamic range of protein abundances, samples were prepared at seven concentrations ranging from 0.1% to 100% human content and processed using the 3 Th with 6 ms maxIT DIA acquisition method, but only 1% to 70% is shown on the plots. The peptides matching the chicken FASTA file were removed from the Skyline document prior to exporting the protein abundance results. The protein-level abundances were normalized to the 100% human reference sample and log₂-transformed (log₂[A/A₁₀₀]) to visualize fold-changes across the dilution series. Both instruments demonstrated excellent linear quantitative behavior across the full dilution range, with clear separation between concentration levels (Figure 6B-C). As expected, protein signals decreased progressively with a lower percentage of human EVs and tracked the expected dilution ratio consistently.

**Figure 6.**
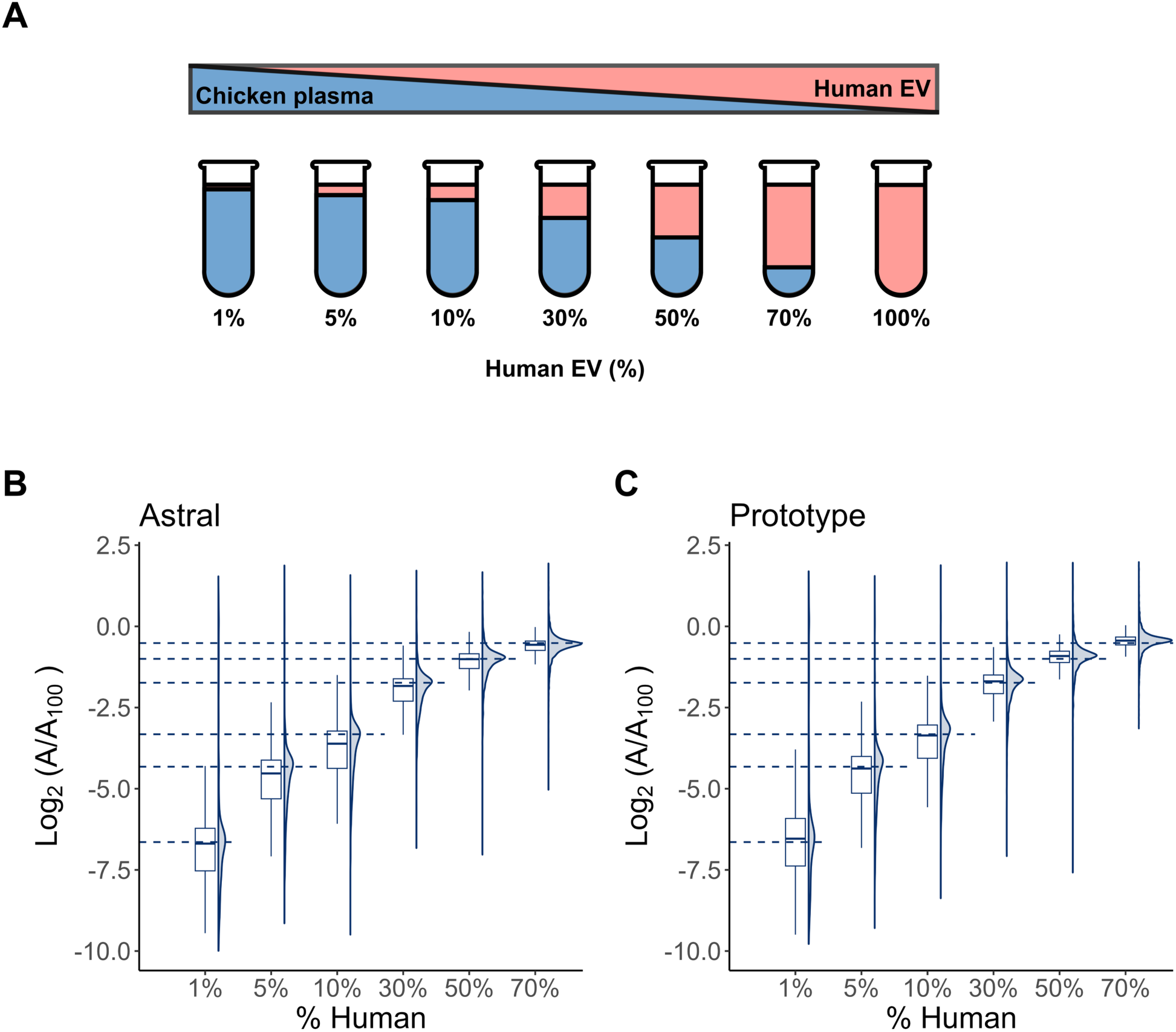
Matrix-matched calibration of human extracellular vesicle (EV) proteins in a chicken plasma background. **(A)** Schematic of volumetric mixing used to generate a dilution series of human extracellular vesicle (EV) peptides in chicken extracellular peptides. The percentage values indicate the volumetric proportion of human EV peptides in each mixture, with the remainder composed of chicken EV peptides. All mixtures were processed using peptide samples generated by Mag-Net^21^. Following a DIA-NN search using a Carafe spectral library, the peptides shared between chicken and human were excluded. Quantitative analysis was performed using Skyline, and protein areas were normalized to those in the 100% human sample to obtain log₂-transformed abundance ratios. Panels show the distribution of log₂(A/A₁₀₀) values for human peptides quantified on the **(B)** Orbitrap Astral MS and **(C)** Orbitrap Astral Zoom MS prototype. Dashed lines denote the expected log₂ ratios based on dilution.

## Discussion

The Orbitrap Astral Zoom MS prototype offers several clear advantages over the standard Orbitrap Astral MS across key performance metrics relevant to quantitative proteomics using DIA. Specifically, the prototype instrument can achieve faster MS2 acquisition rates, reduced acquisition overhead, and shorter cycle times. Importantly, the enhanced acquisition efficiency also enables a valuable trade-off: users can extend injection times to improve ion sampling while still maintaining acquisition rates comparable to the standard Orbitrap Astral MS. For example, with 2 Th isolation windows, the Orbitrap Astral Zoom MS prototype achieves the same acquisition rate at >7 ms injection time as the Orbitrap Astral MS does at 6 ms. This added flexibility, enabled by the pre-accumulation step, makes the Orbitrap Astral Zoom MS prototype well-suited for applications that need both sensitivity and speed, such as low input or single-cell applications.

We benchmarked these two instruments by further evaluating the number of precursor and protein detections, the quantitative precision, ion count, and quantitative across three different DIA methods and input material. Across all conditions of HeLa peptide input, the Orbitrap Astral Zoom MS prototype detected more precursors and proteins than the Orbitrap Astral MS with lower median CVs. One of the more striking findings was the ability of the Orbitrap Astral Zoom MS prototype to accumulate more ions per peptide. This was evident across several ion count metrics and remained true after calibration using instrument-specific α correction factors. The Orbitrap Astral Zoom MS prototype not only achieved AGC targets more effectively but also yielded higher total ion counts across the LC peak, suggesting that more ions contributed to each peptide’s quantification. These benefits were especially evident in peptides near the detection threshold, where modest gains in ion sampling can significantly improve detectability and precision. Despite these differences, the Orbitrap Astral MS and Orbitrap Astral Zoom MS prototype deliver quantitatively precise measurements.

We show here that mass spectrometers can be calibrated using the well-established relationship between the number of ions and measurement variance. This calibration standardizes the signal response to ions per second, enabling meaningful comparisons between different instruments and methods. It also lets us determine how many ions contribute to each measurement even from instruments that originally had different and arbitrary intensity scales.

It is standard for new quantitative methods to report the coefficient of variation (CV) as a measure of precision. However, these CV values are often calculated from a small same size (frequently fewer than 10 replicates), which limits their reliability. Because of the established relationship between the number of ions and variance, we propose that reporting the number of ions used in each measurement provides a valuable complement beyond standard precision figures of merit. When both ion count and precision are reported together, researchers can better assess whether a reported CV might be misleadingly optimistic due to small or biased sampling. This dual reporting approach offers a more complete picture of measurement quality and reliability.

Additionally, both instruments demonstrate strong quantitative accuracy across a dynamic range of peptide inputs and sample types, including matrix-matched human EV samples. While the Orbitrap Astral Zoom MS prototype outperforms the Orbitrap Astral MS in most metrics, the Orbitrap Astral MS remains a robust and reliable platform for DIA-based proteomics.

## Data availability

Instrument raw files, database search results, and Skyline files are available on Panorama at https://panoramaweb.org/MacCoss_ModifiedOrbitrapAstralZoom.url. R code for statistical analysis and plotting is available on GitHub (https://github.com/uw-maccosslab/manuscript-ion-counting).

## Acknowledgements

This work was supported in part by National Institutes of Health grants R24GM141156, U19AG065156, U01DK137097, and T32AG052354. Additionally, support was provided by the Intelligence Advanced Research Projects Activity (IARPA) TEI-REX program under contract numbers W911NF2220059. We thank Drs. Nicholas Riley and Tim Veth from the Riley lab in the Department of Chemistry at the University of Washington for generously providing access to their Orbitrap Ascend mass spectrometer.

## SUPPLEMENTARY MATERIALS

### SUPPLEMENTARY FIGURES

**Supplementary Figure 1.**
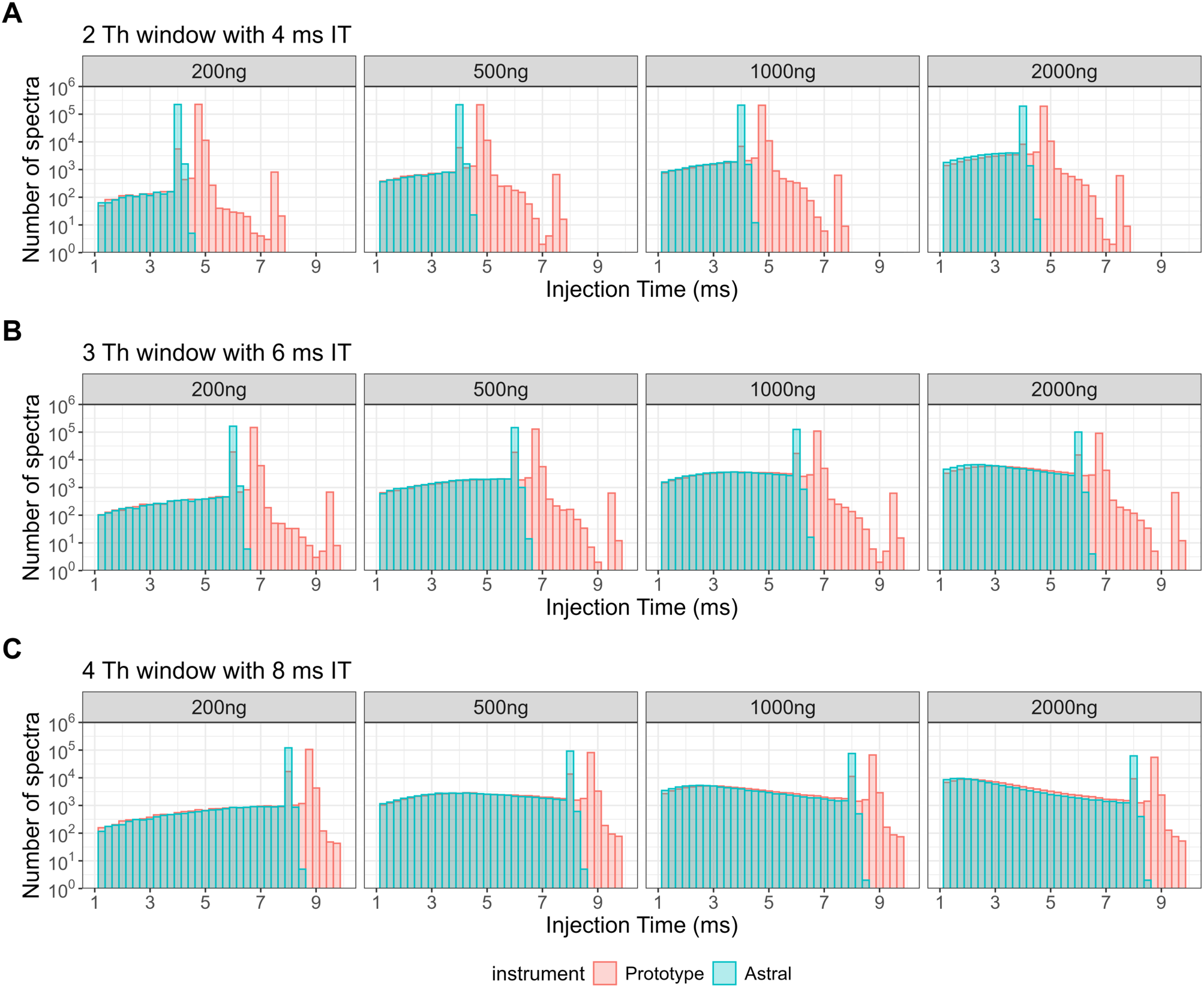
Injection time comparison between different isolation windows and input HeLa masses. Histograms show the number of MS2 spectra acquired at each injection time (IT) for the Orbitrap Astral MS (blue) and Orbitrap Astral Zoom MS prototype (red) across five HeLa peptide input amounts (200–2000 ng) and three DIA acquisition settings: **(A)** 2 Th with 4 ms IT, **(B)** 3 Th with 6 ms IT, and **(C)** 4 Th with 8 ms IT.

**Supplementary Table 1.**
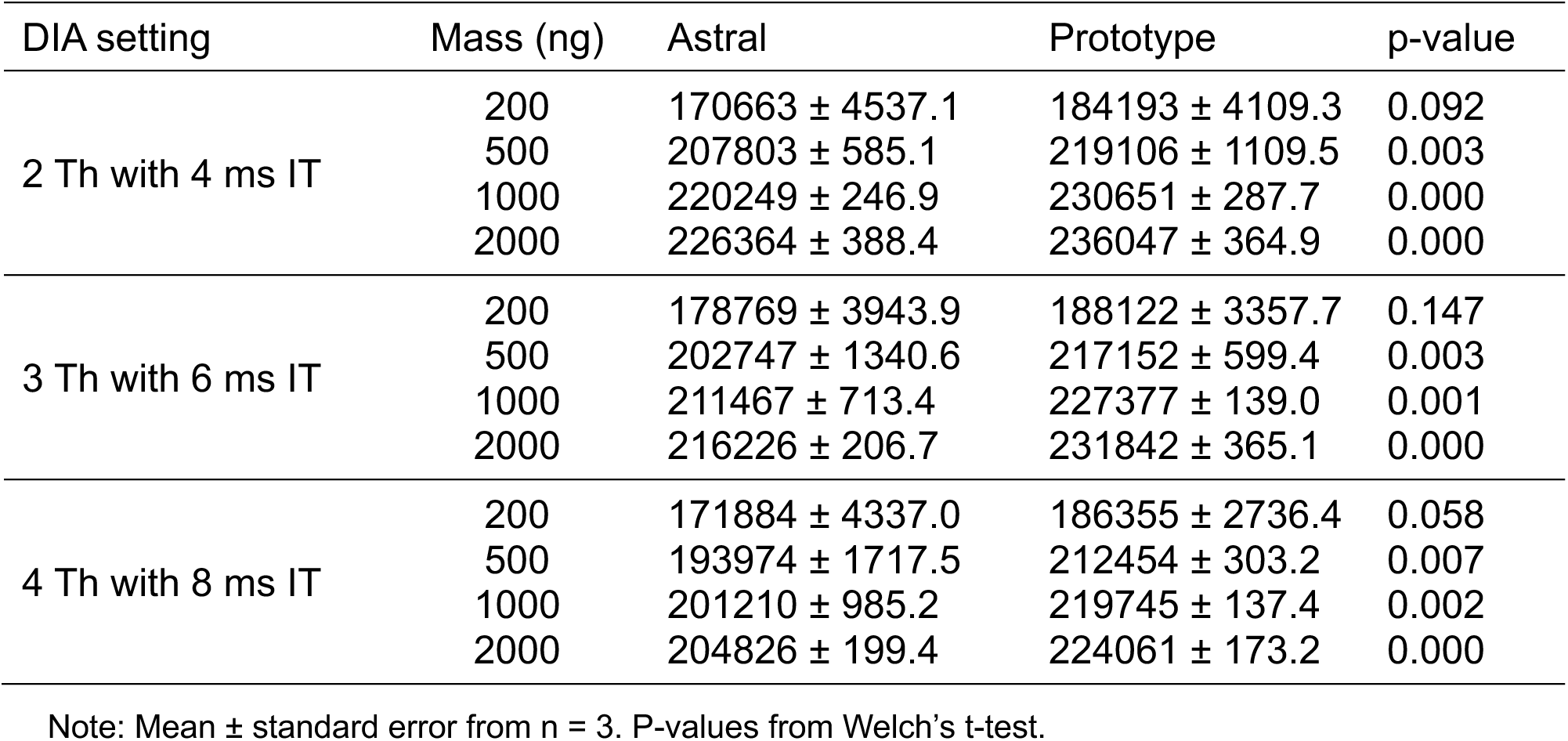
Precursors detected by DIA-NN search with reported p-values.

**Supplementary Table 2.**
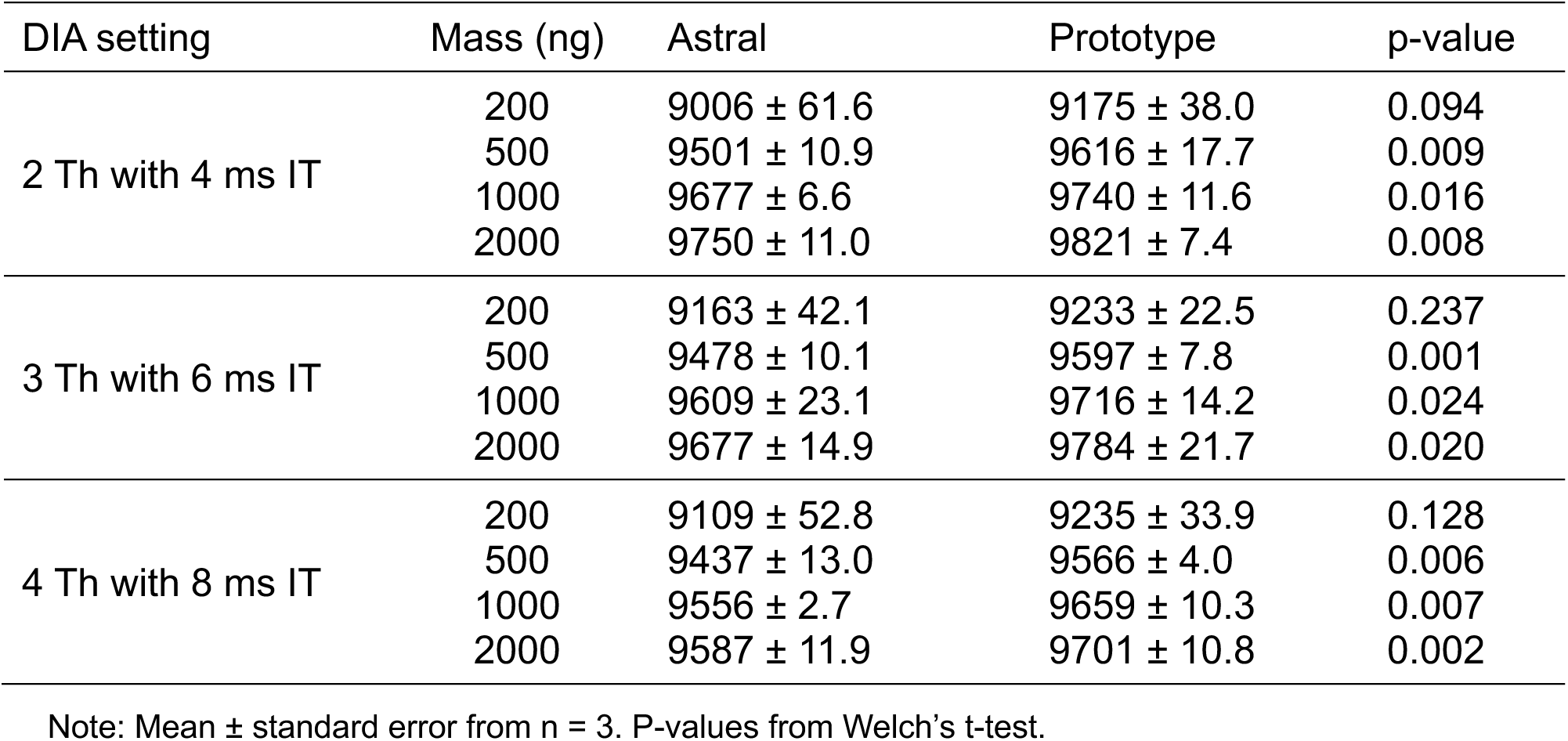
Proteins detected by DIA-NN search with reported p-values.

**Supplementary Figure 2.**
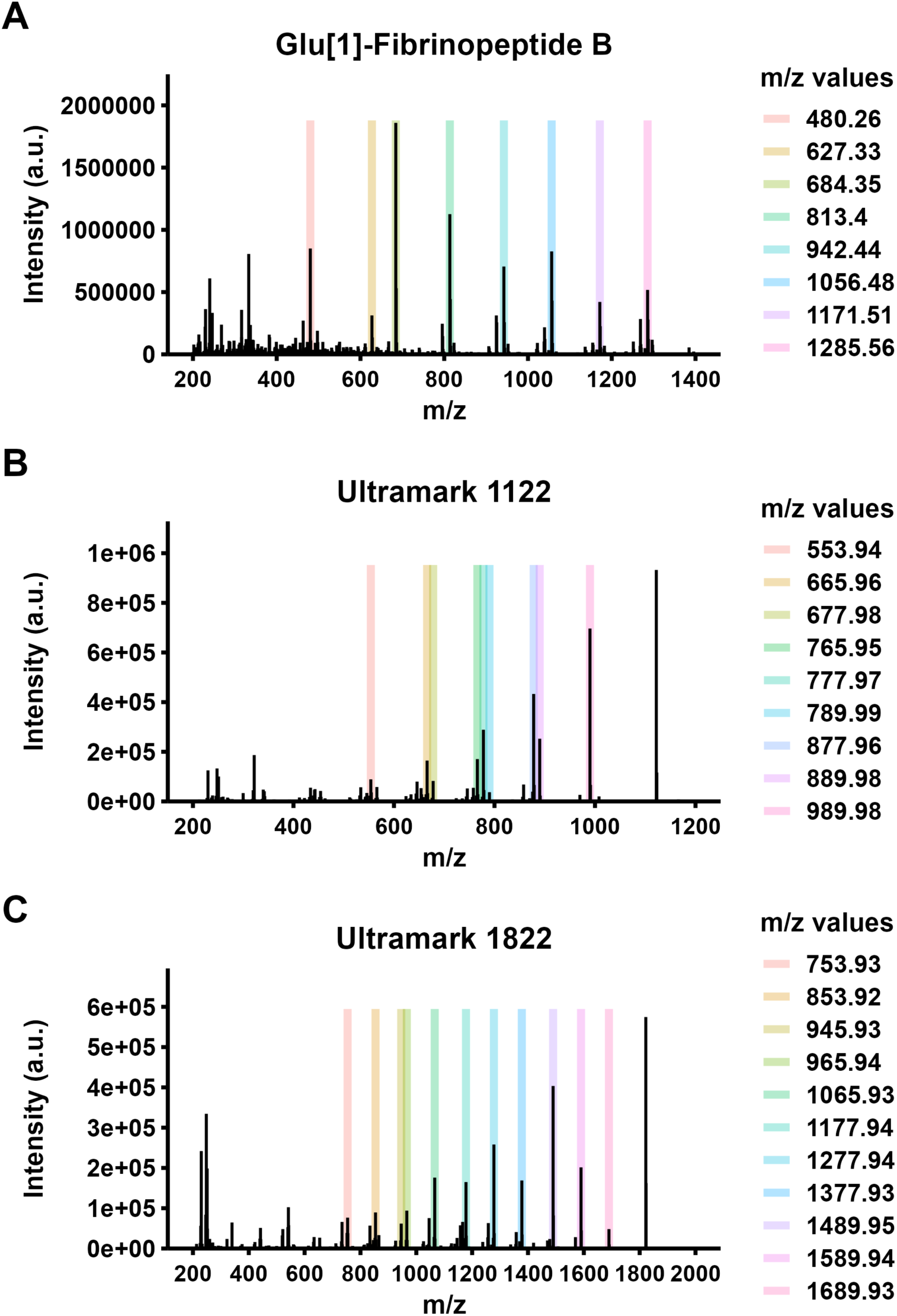
Reference compound MS2 spectra used for ion intensity calibration. Representative tandem mass spectra of three calibration standards used for cross-instrument intensity calibration from Orbitrap Astral MS: **(A)** Glu[1]-Fibrinopeptide B, **(B)** Ultramark 1122, and **(C)** Ultramark 1822. Highlighted colored bars indicate selected fragment m/z peaks used in the calibration analysis. These fragment ions were used to construct calibration curves correlating observed intensity and variance to ion count, enabling conversion of instrument-reported intensities to ions per second (ions/sec). m/z values for each set of reference ions are listed to the right of each panel.

**Supplementary Figure 3.**
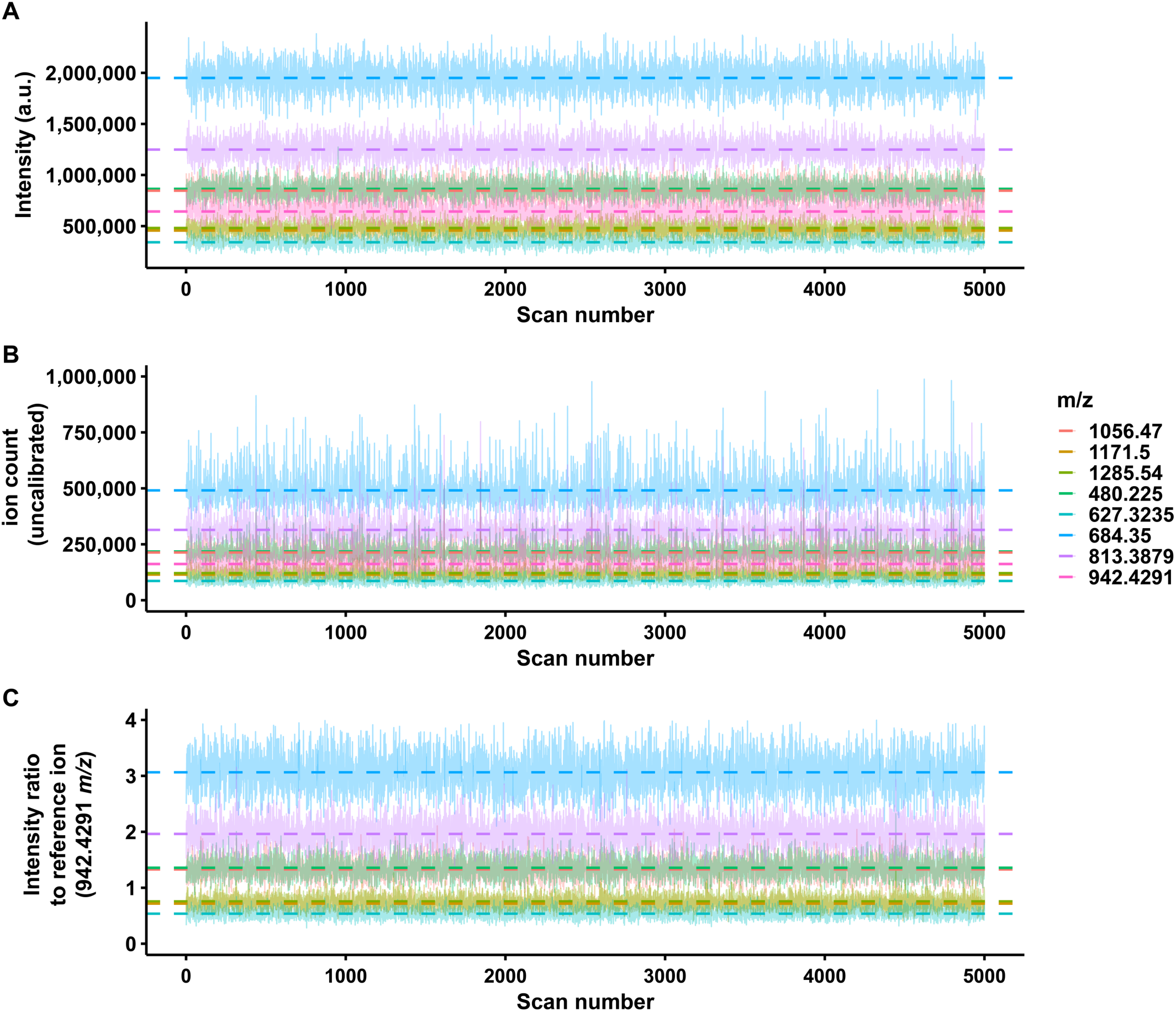
Glu[1]-Fibrinopeptide B fragment ion intensities, uncalibrated ion counts, and intensity ratios across replicate MS2 spectra. Eight fragment ions (colored by *m/z*) from Glu[1]-Fibrinopeptide B were monitored across 5000 MS2 spectra from the Orbitrap Astral mass spectrometer. The dashed lines indicate the median value across all plots. (A) Raw intensities in arbitrary units of each fragment ion. (B) Uncalibrated ion counts computed as the product of intensity and injection time. (C) Intensity ratios of each fragment ion to the reference ion (942.4291 *m/z*). The intensity ratio is used to assess the measured variance and calculate the mean ratio for ion calibration.

**Supplementary Figure 4.**
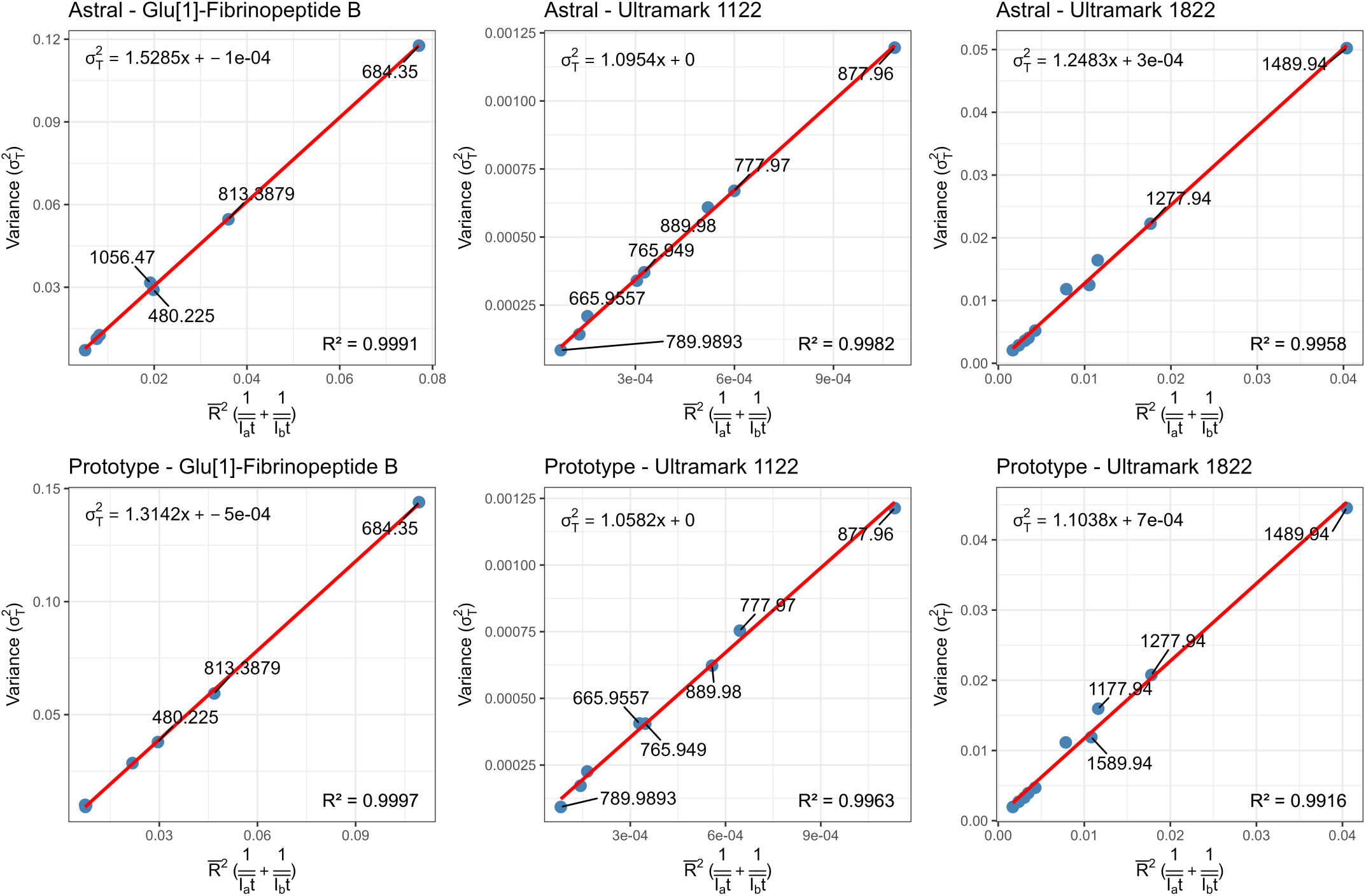
Calibration of measured MS2 intensity to ions per second for the Orbitrap Astral MS and Orbitrap Astral Zoom MS prototype instruments. Linear regression models were generated to calibrate MS2 ion intensity to absolute ion counts using variance-based estimates from Poisson statistics. Each plot shows the squared measurement standard deviation (σ^2^ₜ) as a function of the mean ratio signal intensity and number of ions (x-axis) for selected fragment ions from Glu[1]-Fibrinopeptide B, Ultramark 1122, and Ultramark 1822. Top row shows calibration results for the Orbitrap Astral MS; bottom row for the Orbitrap Astral Zoom MS prototype. The slope of the linear regression line represents the α (alpha) factor. R^2^ values indicate model goodness-of-fit. Each labeled point corresponds to a distinct m/z fragment ion used for calibration.

**Supplementary Table 3.**
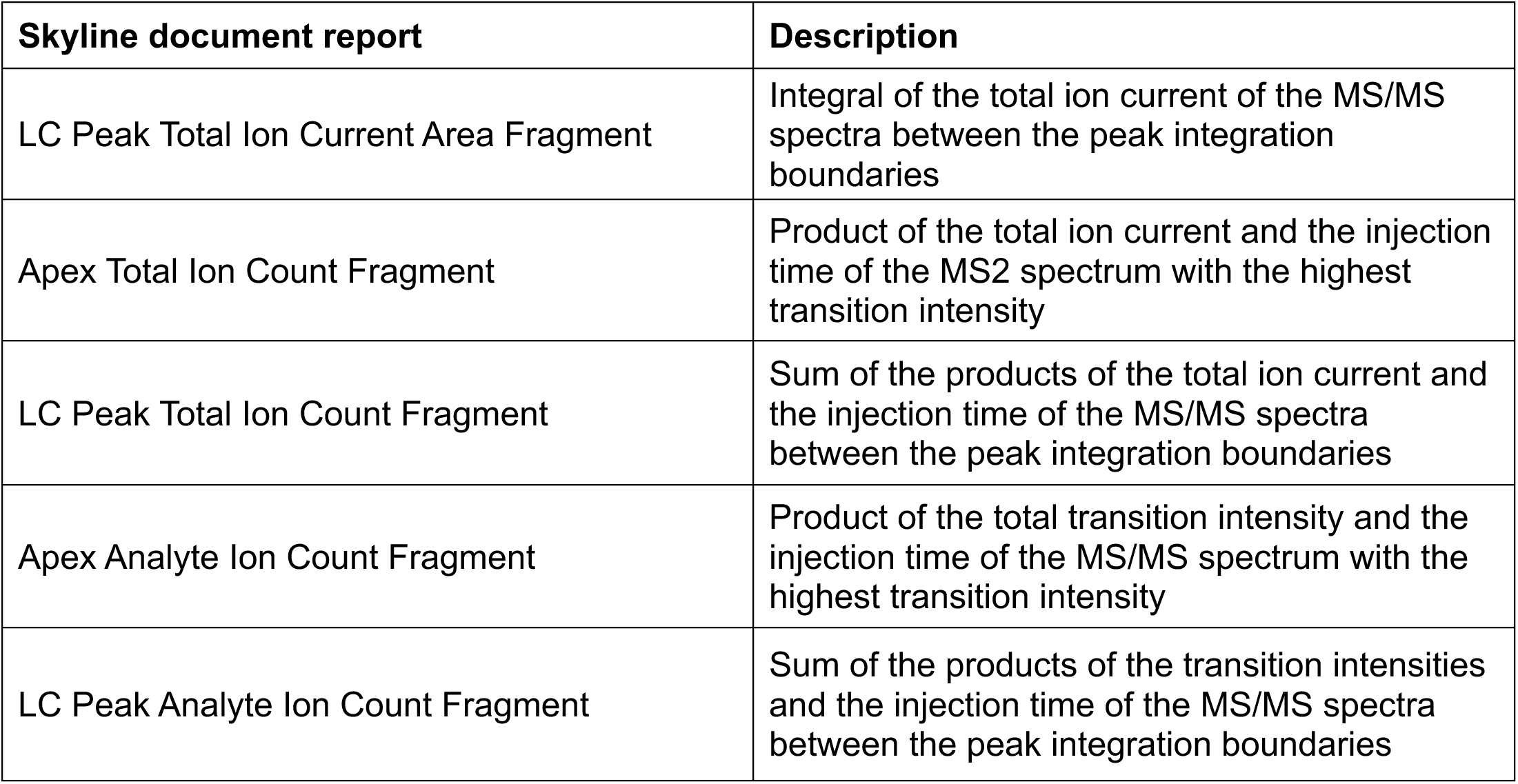
Skyline document reports outlining the new ion counting features.

**Supplementary Figure 5.**
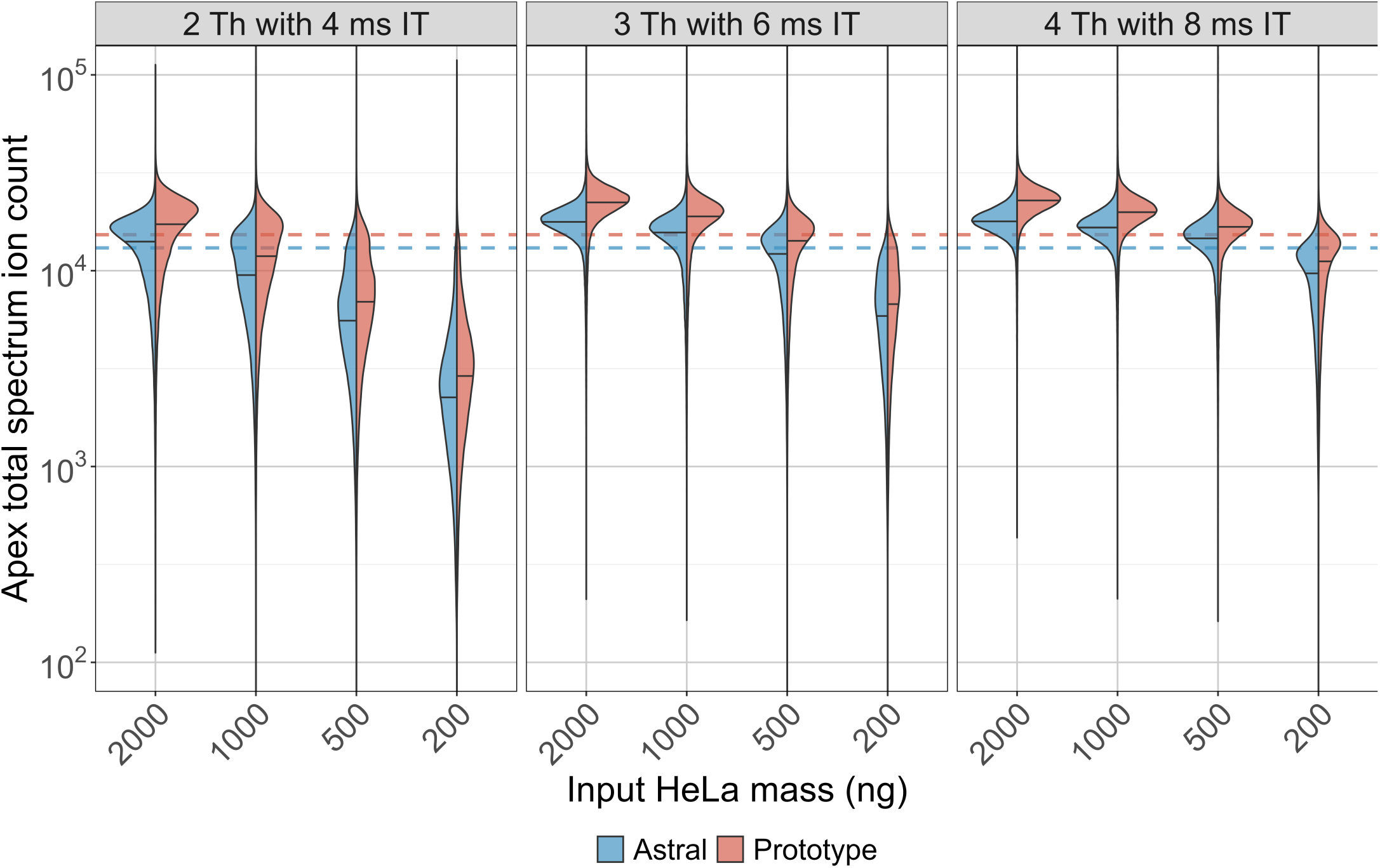
Apex total spectrum ion counts for Orbitrap Astral MS and Orbitrap Astral Zoom MS prototype instruments across input amounts and DIA settings. Violin plots show the distribution of apex total ion counts (in ions/sec) across HeLa peptide inputs ranging from 200 to 2000 ng for three DIA acquisition settings: 2 Th with 4 ms IT, 3 Th with 6 ms IT, and 4 Th with 8 ms IT. Ion counts were calibrated to ions/sec using instrument-specific α correction factors (1.53 for Orbitrap Astral MS and 1.31 for Orbitrap Astral Zoom MS prototype). The apex ion count was calculated as the product of MS2 injection time and total ion current (TIC) at the spectrum with the highest transition intensity for each peptide. Dashed horizontal lines indicate the calibrated target AGC value of 20,000 ions for each instrument.

**Supplementary Figure 6.**
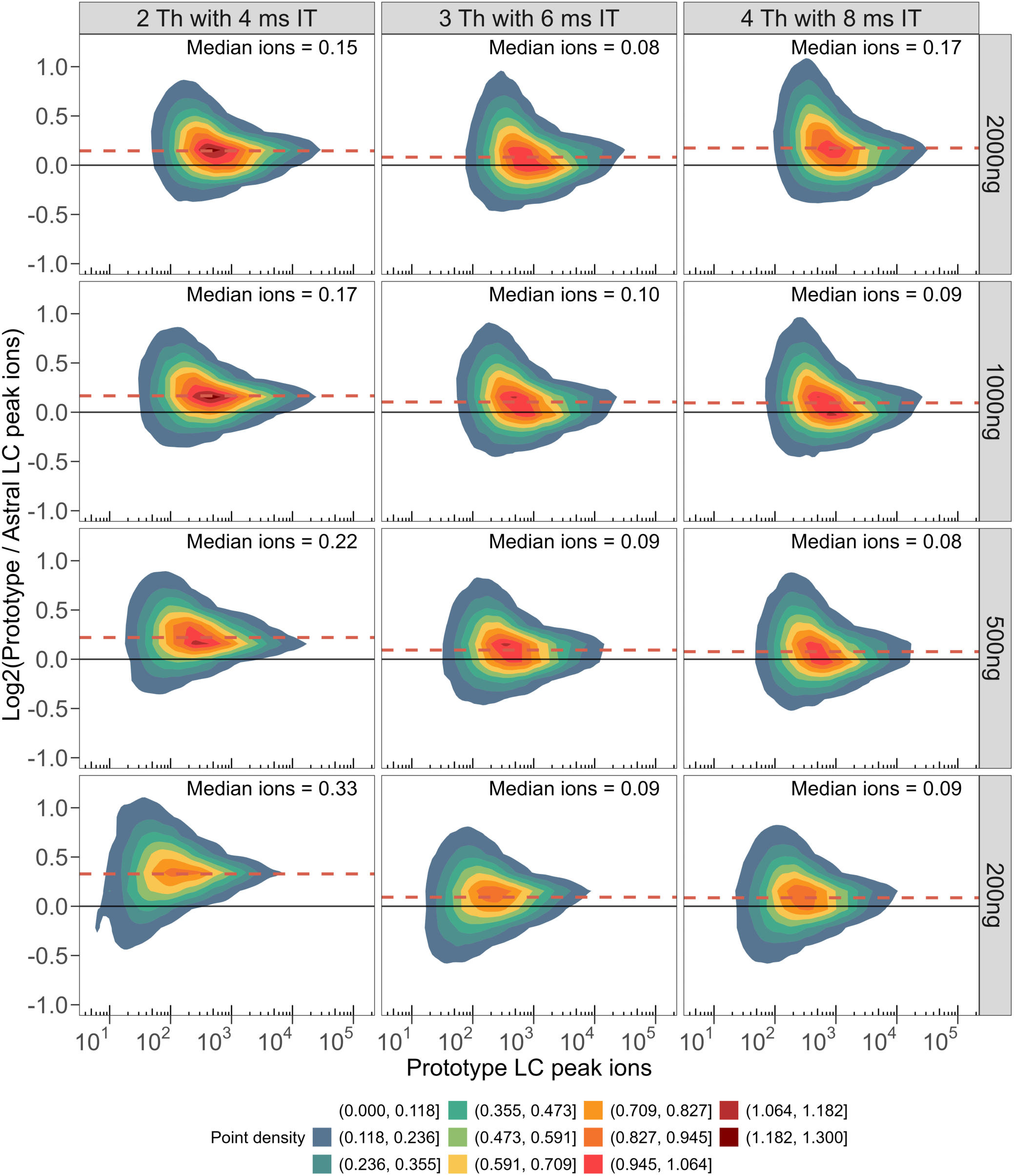
Full distribution of the paired peptide ratio between Orbitrap Astral MS and Orbitrap Astral Zoom MS prototype LC peak peptide ions. Each subplot shows a 2D kernel density estimate comparing the log₂ ratio of LC peak ion intensities (Orbitrap Astral Zoom MS prototype / Orbitrap Astral MS) with the Orbitrap Astral Zoom MS prototype LC peak ion intensity on the x-axis. Rows represent different sample loads (2000 to 200 ng), while columns reflect different DIA acquisition settings—2 Th with 4 ms IT, 3 Th with 6 ms IT, and 4 Th with 8 ms IT. The solid black horizontal line indicates no change (log₂ ratio = 0), while the dashed red line denotes the median log₂ fold change.

**Supplementary Figure 7.**
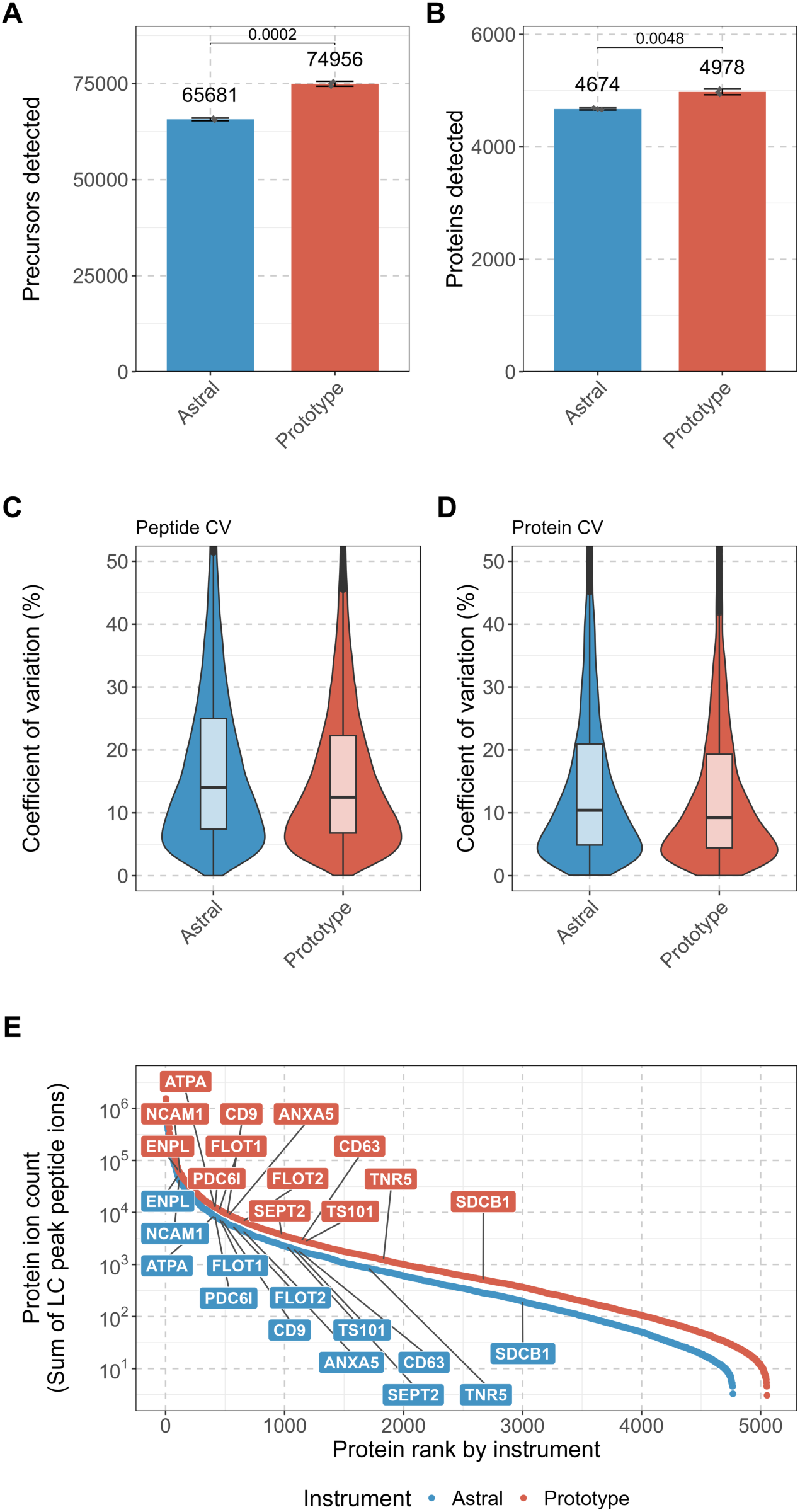
Comparison of performance metrics between the Orbitrap Astral MS and Orbitrap Astral Zoom MS prototype for the human extracellular vesicles (EV). Total number of (A) precursors detected and (B) proteins detected by DIA-NN 2.1.0 search with p-values (unpaired two-tailed t-tests) shown above each bar. The calculated coefficient of variation between the (C) peptide-abundances and (D) protein abundances. (E) Ranked protein abundance curves of summed LC peak peptide ion counts per protein with select EV markers proteins labeled to highlight differences in ion sampling depth between instruments.

