## Supplementary materials for "Evaluation of a prototype Orbitrap Astral Zoom mass spectrometer for quantitative proteomics – Beyond identification lists"


### SUPPLEMENTARY FIGURES:


- Figure S1. Injection time comparison between different isolation windows and input HeLa masses.
- Table S1. Precursors detected by DIA-NN search with reported p-values.
- Table S2. Proteins detected by DIA-NN search with reported p-values.
- Figure S2. Reference compound MS2 spectra used for ion calibration.
- Figure S3. Glu[1]-Fibrinopeptide B fragment ion intensities, uncalibrated ion counts, and intensity ratios across replicate MS2 spectra
- Figure S4. Calibration of measured MS2 intensity to ions per second for the Orbitrap Astral MS and Orbitrap Astral Zoom MS prototype instruments.
- Table S3. Skyline document reports outlining the new ion counting features.
- Figure S5. Apex total spectrum ion counts for Orbitrap Astral MS and Orbitrap Astral Zoom MS prototype instruments across input amounts and DIA settings.
- Figure S6. Full distribution of the paired peptide ratio between Orbitrap Astral MS and Orbitrap Astral Zoom MS prototype LC peak peptide ions.
- Figure S7. Comparison of performance metrics between the Orbitrap Astral MS and Orbitrap Astral Zoom MS prototype for the human extracellular vesicles (EV).

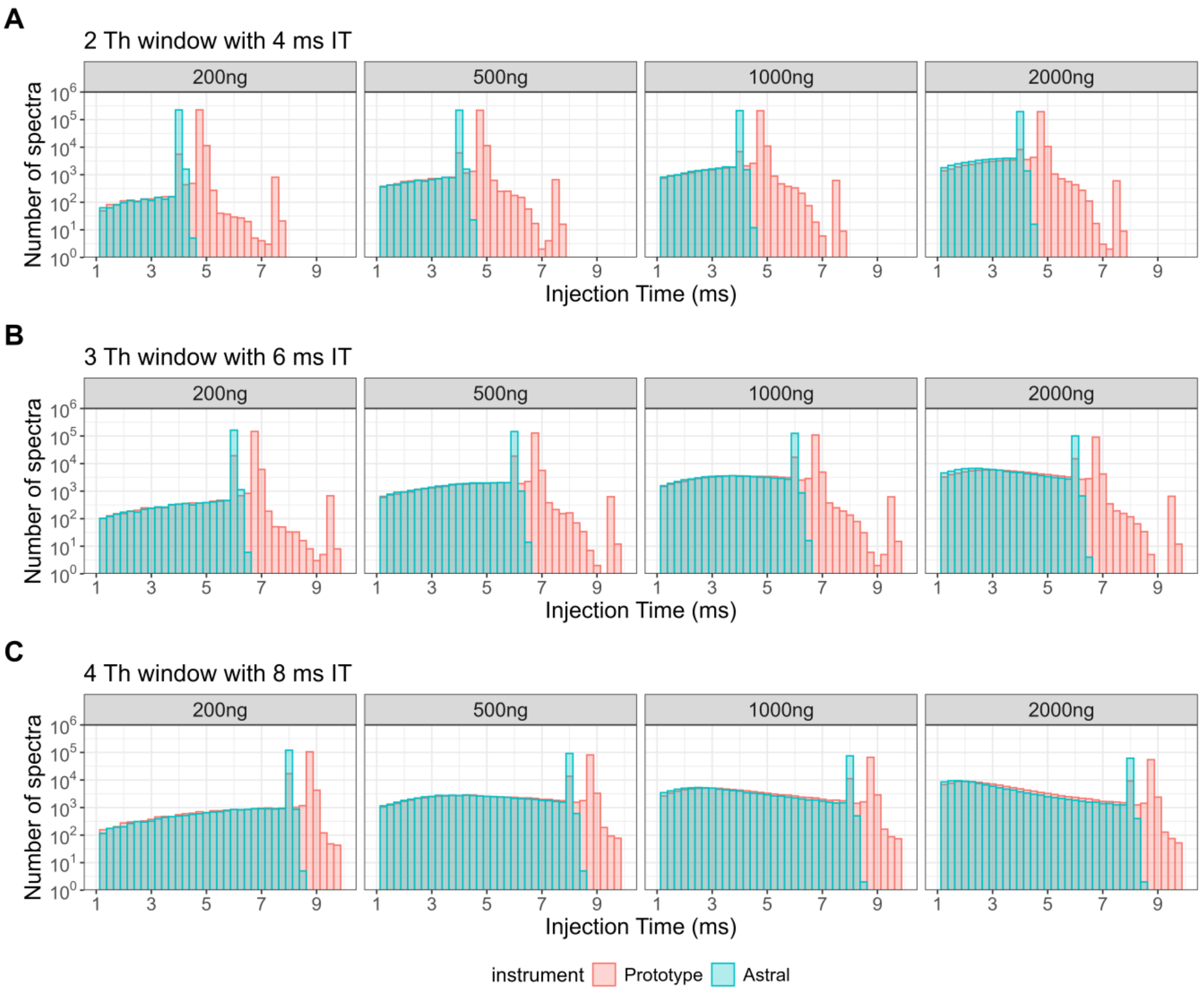


**Supplementary Figure 1. Injection time comparison between different isolation windows and input HeLa masses.** Histograms show the number of MS2 spectra acquired at each injection time (IT) for the Orbitrap Astral MS (blue) and Orbitrap Astral Zoom MS prototype (red) across five HeLa peptide input amounts (200–2000 ng) and three DIA acquisition settings: **(A)** 2 Th with 4 ms IT, **(B)** 3 Th with 6 ms IT, and **(C)** 4 Th with 8 ms IT.

**Supplementary Table 1.** Precursors detected by DIA-NN search with reported p-values

| DIA setting | Mass (ng) | Astral | Prototype | p-value |
|---|---|---|---|---|
| 2 Th with 4 ms IT | 200 | 170663 ± 4537.1 | 184193 ± 4109.3 | 0.092 |
| | 500 | 207803 ± 585.1 | 219106 ± 1109.5 | 0.003 |
| | 1000 | 220249 ± 246.9 | 230651 ± 287.7 | 0.000 |
| | 2000 | 226364 ± 388.4 | 236047 ± 364.9 | 0.000 |
| 3 Th with 6 ms IT | 200 | 178769 ± 3943.9 | 188122 ± 3357.7 | 0.147 |
| | 500 | 202747 ± 1340.6 | 217152 ± 599.4 | 0.003 |
| | 1000 | 211467 ± 713.4 | 227377 ± 139.0 | 0.001 |
| | 2000 | 216226 ± 206.7 | 231842 ± 365.1 | 0.000 |
| 4 Th with 8 ms IT | 200 | 171884 ± 4337.0 | 186355 ± 2736.4 | 0.058 |
| | 500 | 193974 ± 1717.5 | 212454 ± 303.2 | 0.007 |
| | 1000 | 201210 ± 985.2 | 219745 ± 137.4 | 0.002 |
| | 2000 | 204826 ± 199.4 | 224061 ± 173.2 | 0.000 |

Note: Mean ± standard error from n = 3. P-values from Welch’s t-test.

**Supplementary Table 2.** Proteins detected by DIA-NN search with reported p-values

| DIA setting | Mass (ng) | Astral | Prototype | p-value |
|---|---|---|---|---|
| 2 Th with 4 ms IT | 200 | 9006 ± 61.6 | 9175 ± 38.0 | 0.094 |
| | 500 | 9501 ± 10.9 | 9616 ± 17.7 | 0.009 |
| | 1000 | 9677 ± 6.6 | 9740 ± 11.6 | 0.016 |
| | 2000 | 9750 ± 11.0 | 9821 ± 7.4 | 0.008 |
| 3 Th with 6 ms IT | 200 | 9163 ± 42.1 | 9233 ± 22.5 | 0.237 |
| | 500 | 9478 ± 10.1 | 9597 ± 7.8 | 0.001 |
| | 1000 | 9609 ± 23.1 | 9716 ± 14.2 | 0.024 |
| | 2000 | 9677 ± 14.9 | 9784 ± 21.7 | 0.020 |
| 4 Th with 8 ms IT | 200 | 9109 ± 52.8 | 9235 ± 33.9 | 0.128 |
| | 500 | 9437 ± 13.0 | 9566 ± 4.0 | 0.006 |
| | 1000 | 9556 ± 2.7 | 9659 ± 10.3 | 0.007 |
| | 2000 | 9587 ± 11.9 | 9701 ± 10.8 | 0.002 |

Note: Mean ± standard error from n = 3. P-values from Welch’s t-test.

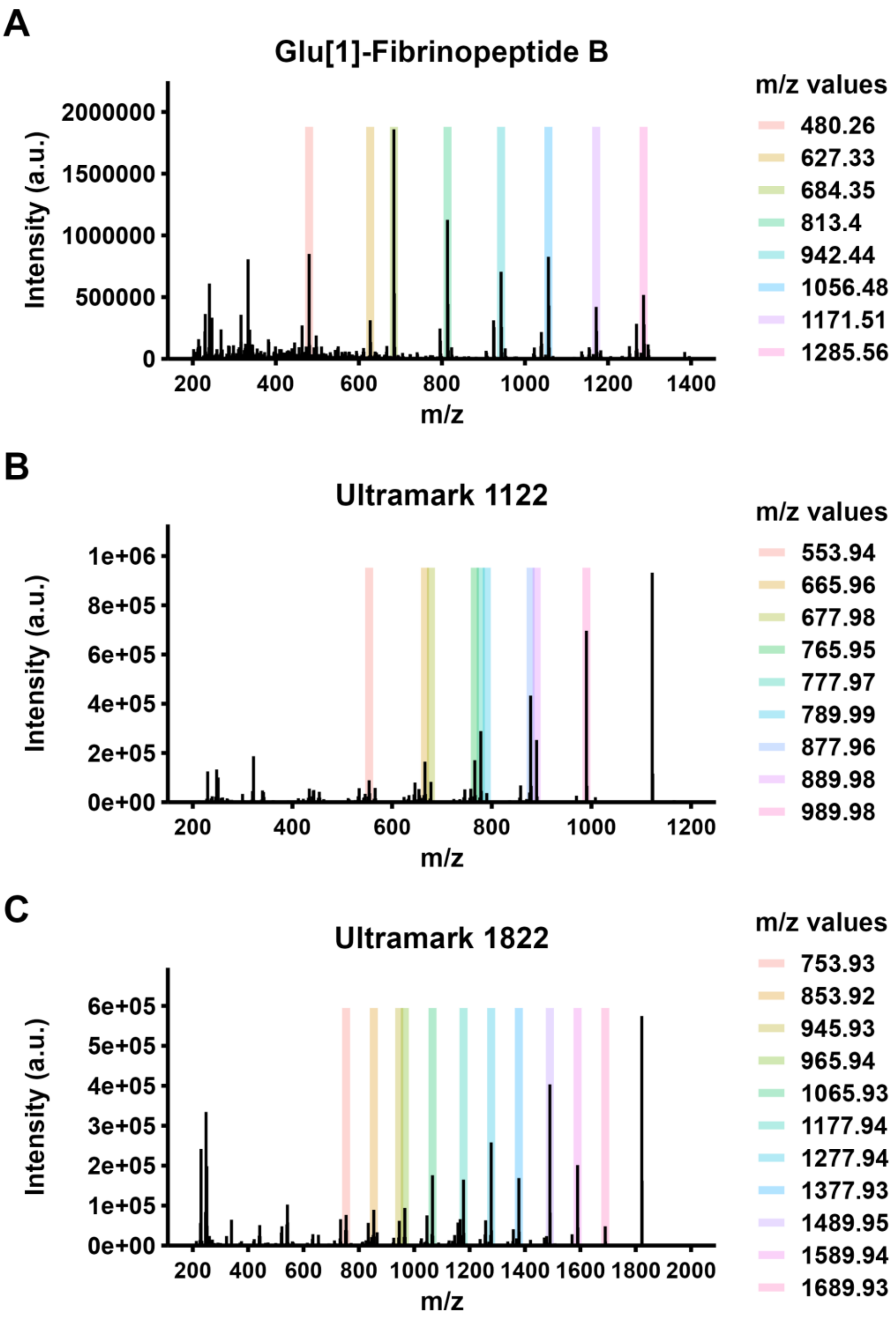


**Supplementary Figure 2. Reference compound MS2 spectra used for ion intensity calibration.** Representative tandem mass spectra of three calibration standards used for cross-instrument intensity calibration from Orbitrap Astral MS: **(A)** Glu[1]-Fibrinopeptide B, **(B)** Ultramark 1122, and **(C)** Ultramark 1822. Highlighted colored bars indicate selected fragment m/z peaks used in the calibration analysis. These fragment ions were used to construct calibration curves correlating observed intensity and variance to ion count, enabling conversion of instrument-reported intensities to ions per second (ions/sec). m/z values for each set of reference ions are listed to the right of each panel.

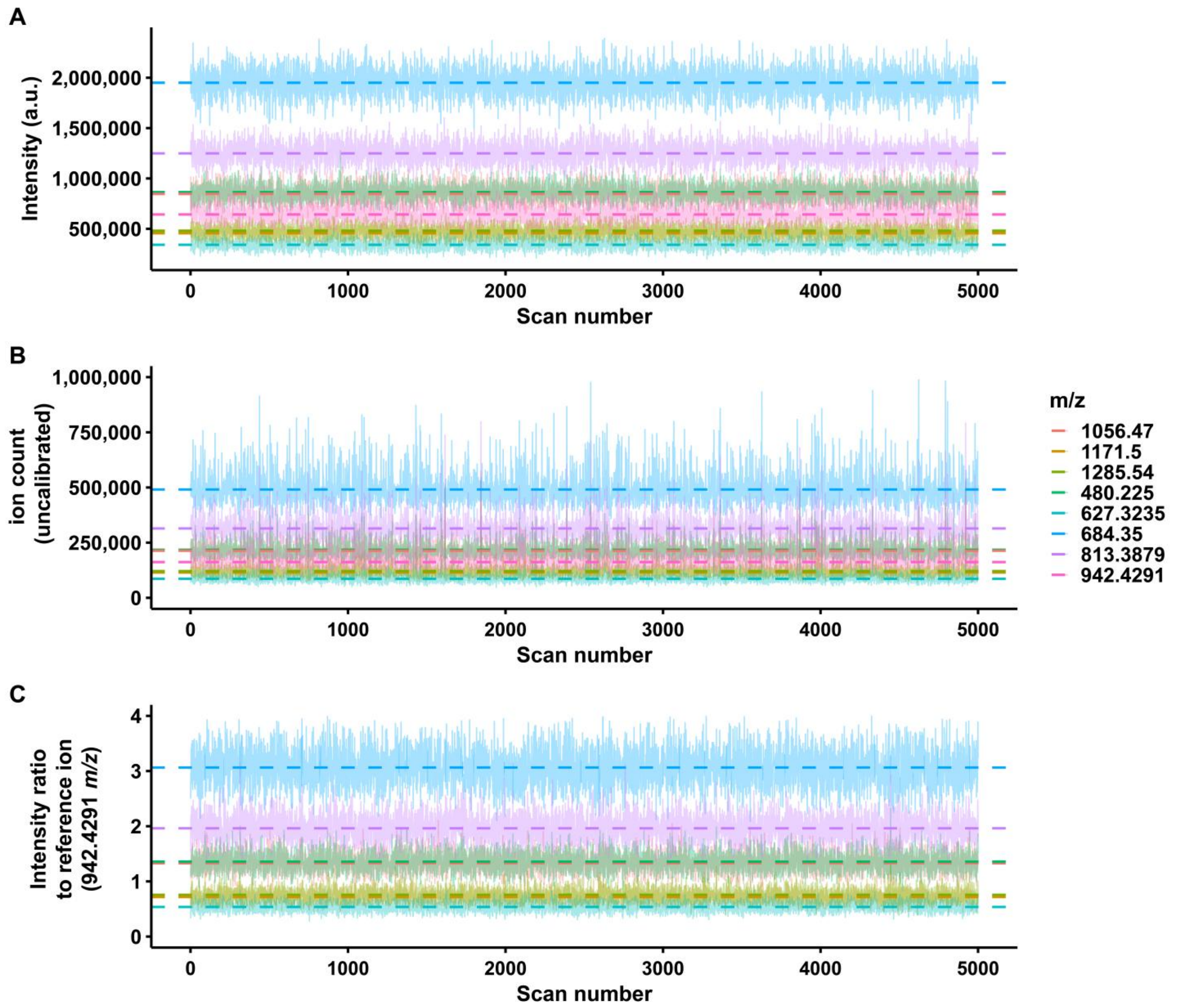


**Supplementary Figure 3.** Glu[1]-Fibrinopeptide B fragment ion intensities, uncalibrated ion counts, and intensity ratios across replicate MS2 spectra. Eight fragment ions (colored by *m/z*) from Glu[1]-Fibrinopeptide B were monitored across 5000 MS2 spectra from the Orbitrap Astral mass spectrometer. The dashed lines indicate the median value across all plots. (A) Raw intensities in arbitrary units of each fragment ion. (B) Uncalibrated ion counts computed as the product of intensity and injection time. (C) Intensity ratios of each fragment ion to the reference ion (942.4291 *m/z*). The intensity ratio is used to assess the measured variance and calculate the mean ratio for ion calibration.

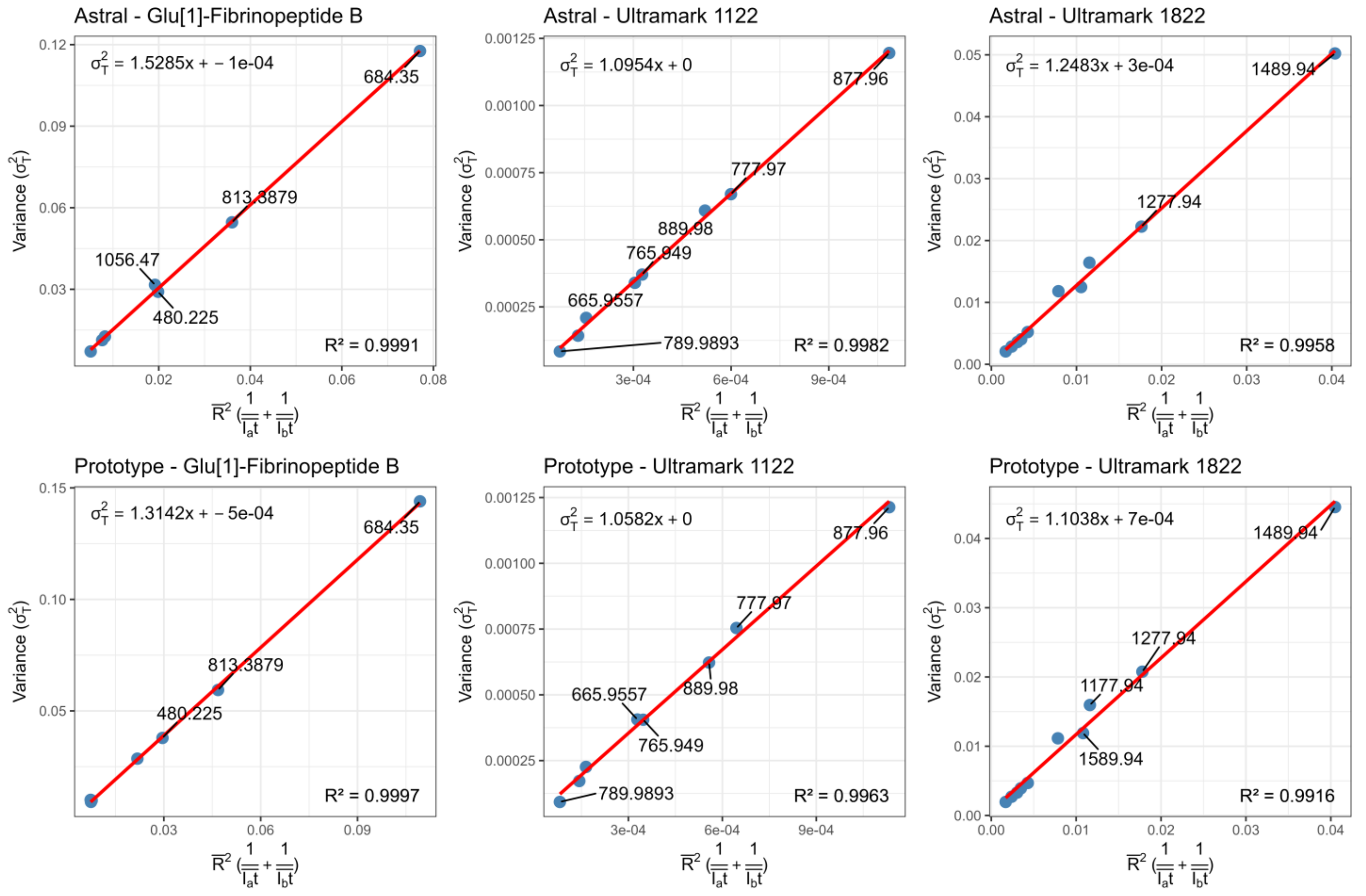


**Supplementary Figure 4.** Calibration of measured MS2 intensity to ions per second for the Orbitrap Astral MS and Orbitrap Astral Zoom MS prototype instruments. Linear regression models were generated to calibrate MS2 ion intensity to absolute ion counts using variance-based estimates from Poisson statistics. Each plot shows the squared measurement standard deviation ($\sigma^2_t$) as a function of the mean ratio signal intensity and number of ions (x-axis) for selected fragment ions from Glu[1]-Fibrinopeptide B, Ultramark 1122, and Ultramark 1822. Top row shows calibration results for the Orbitrap Astral MS; bottom row for the Orbitrap Astral Zoom MS prototype. The slope of the linear regression line represents the α (alpha) factor. $R^2$ values indicate model goodness-of-fit. Each labeled point corresponds to a distinct m/z fragment ion used for calibration.

**Supplementary Table 3**. Skyline document reports outlining the new ion counting features

| **Skyline document report** | **Description** |
|---|---|
| LC Peak Total Ion Current Area Fragment | Integral of the total ion current of the MS/MS spectra between the peak integration boundaries |
| Apex Total Ion Count Fragment | Product of the total ion current and the injection time of the MS2 spectrum with the highest transition intensity |
| LC Peak Total Ion Count Fragment | Sum of the products of the total ion current and the injection time of the MS/MS spectra between the peak integration boundaries |
| Apex Analyte Ion Count Fragment | Product of the total transition intensity and the injection time of the MS/MS spectrum with the highest transition intensity |
| LC Peak Analyte Ion Count Fragment | Sum of the products of the transition intensities and the injection time of the MS/MS spectra between the peak integration boundaries |

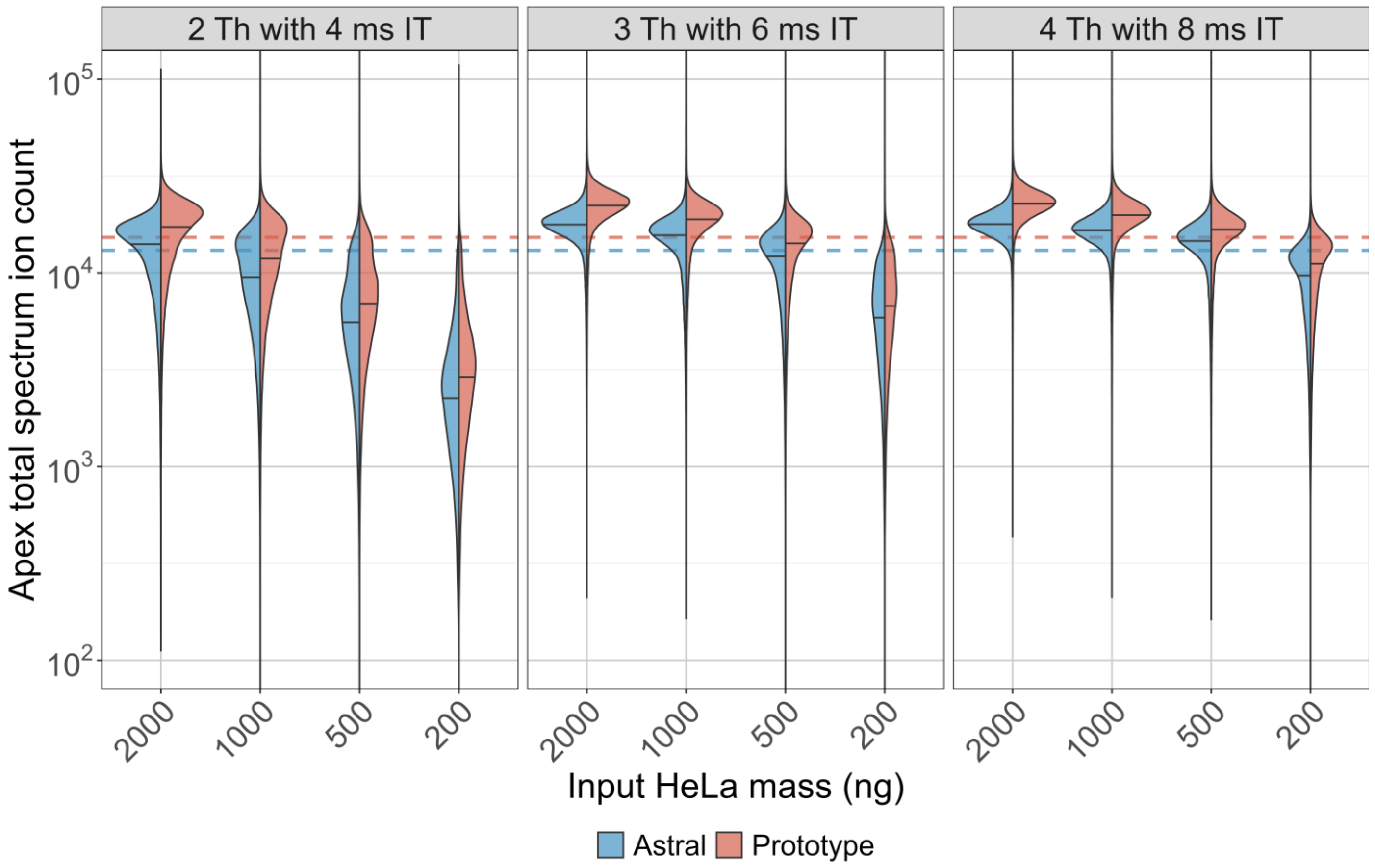


**Supplementary Figure 5.** Apex total spectrum ion counts for Orbitrap Astral MS and Orbitrap Astral Zoom MS prototype instruments across input amounts and DIA settings. Violin plots show the distribution of apex total ion counts (in ions/sec) across HeLa peptide inputs ranging from 200 to 2000 ng for three DIA acquisition settings: 2 Th with 4 ms IT, 3 Th with 6 ms IT, and 4 Th with 8 ms IT. Ion counts were calibrated to ions/sec using instrument-specific α correction factors (1.53 for Orbitrap Astral MS and 1.31 for Orbitrap Astral Zoom MS prototype). The apex ion count was calculated as the product of MS2 injection time and total ion current (TIC) at the spectrum with the highest transition intensity for each peptide. Dashed horizontal lines indicate the calibrated target AGC value of 20,000 ions for each instrument.

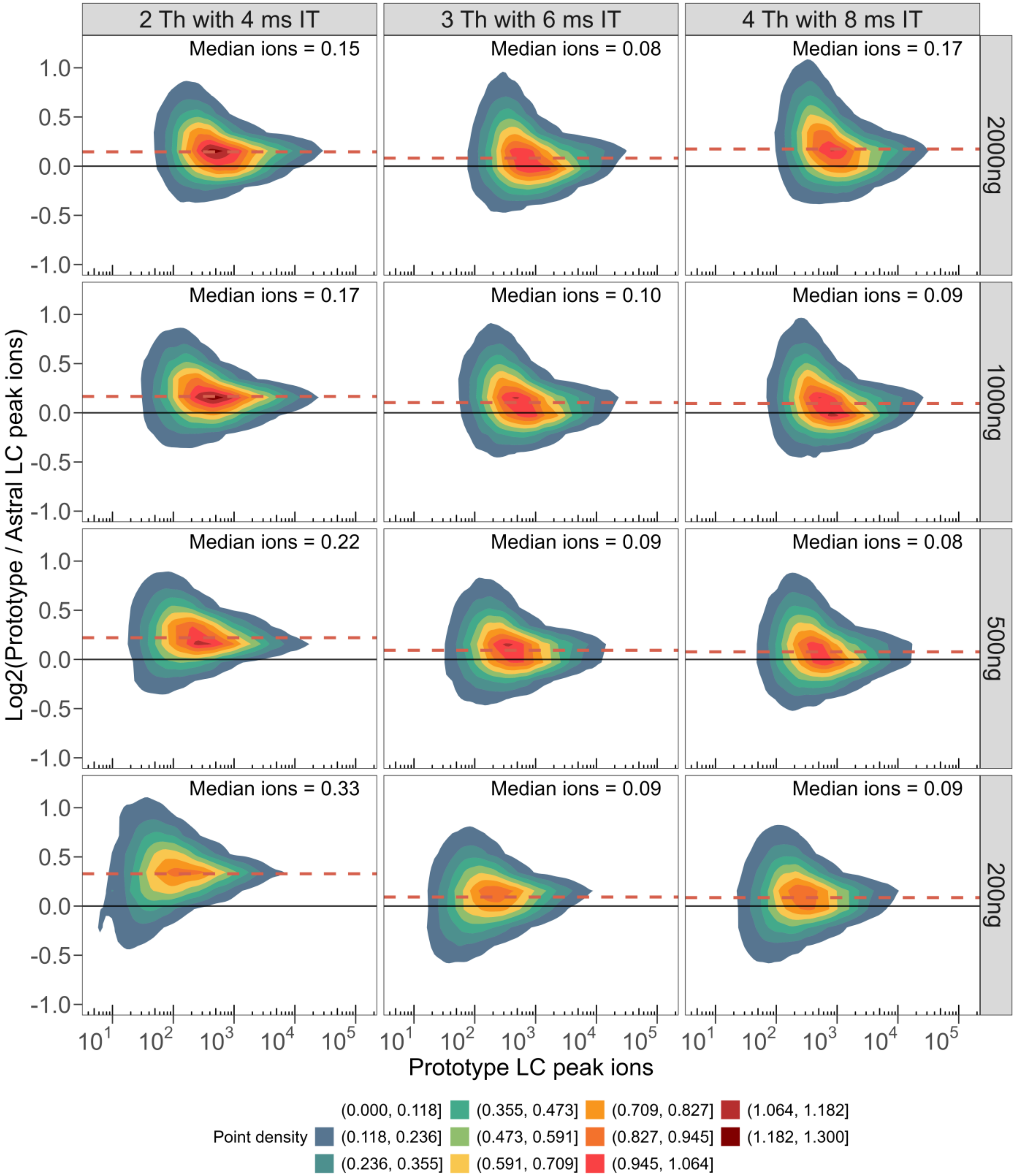


**Supplementary Figure 6.** Full distribution of the paired peptide ratio between Orbitrap Astral MS and Orbitrap Astral Zoom MS prototype LC peak peptide ions. Each subplot shows a 2D kernel density estimate comparing the $\log_2$ ratio of LC peak ion intensities (Orbitrap Astral Zoom MS prototype / Orbitrap Astral MS) with the Orbitrap Astral Zoom MS prototype LC peak ion intensity on the x-axis. Rows represent different sample loads (2000 to 200 ng), while columns reflect different DIA acquisition settings—2 Th with 4 ms IT, 3 Th with 6 ms IT, and 4 Th with 8 ms IT. The solid black horizontal line indicates no change ($\log_2$ ratio = 0), while the dashed red line denotes the median $\log_2$ fold change.

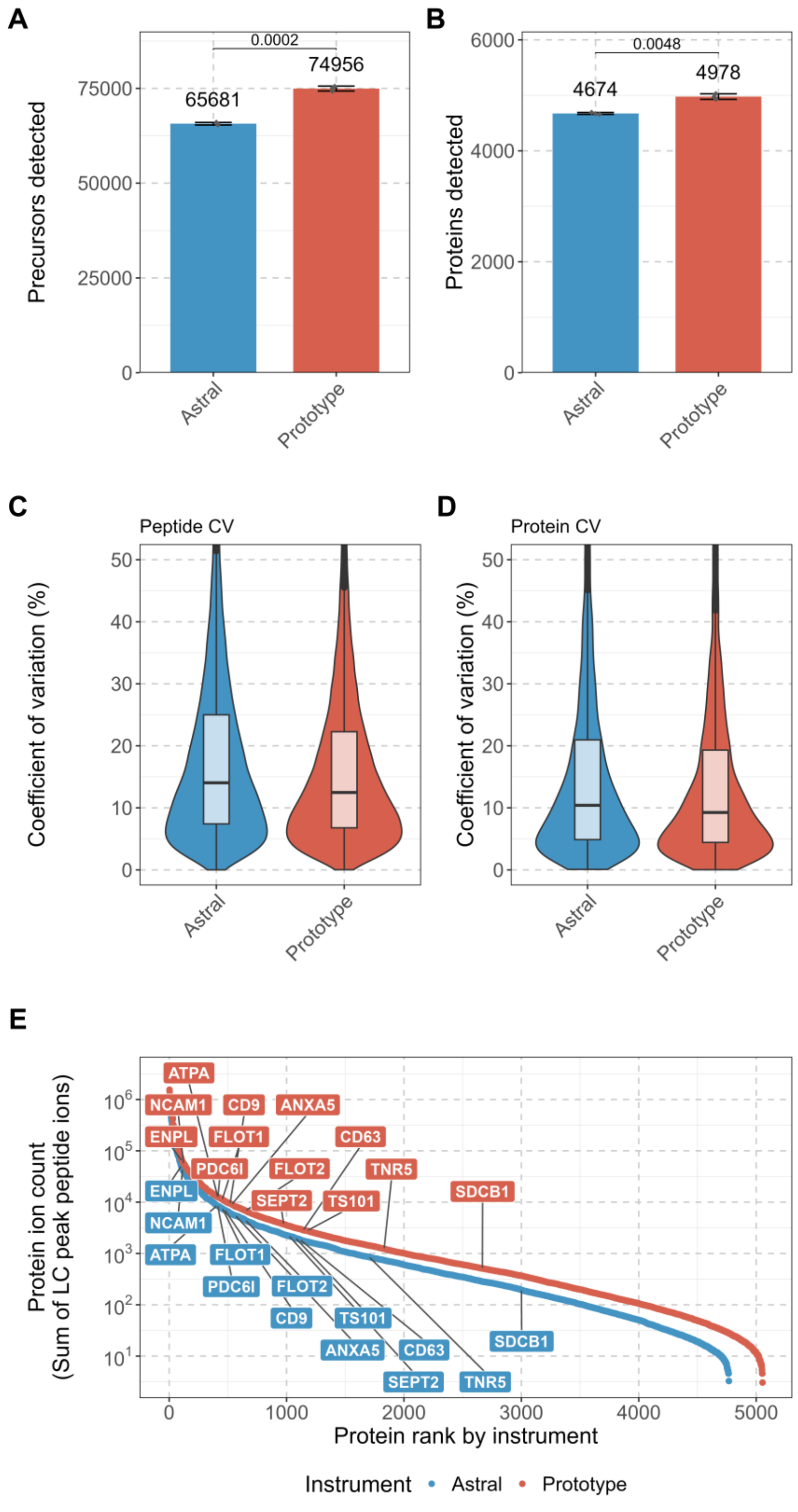


**Supplementary Figure 7.** Comparison of performance metrics between the Orbitrap Astral MS and Orbitrap Astral Zoom MS prototype for the human extracellular vesicles (EV). Total number of (A) precursors detected and (B) proteins detected by DIA-NN 2.1.0 search with p-values (unpaired two-tailed t-tests) shown above each bar. The calculated coefficient of variation between the (C) peptide-abundances and (D) protein abundances. (E) Ranked protein abundance curves of summed LC peak peptide ion counts per protein with select EV markers proteins labeled to highlight differences in ion sampling depth between instruments.
